# Brain Functional Connectome Defines a Transdiagnostic Dimension Shared by Cognitive Dysfunction and Psychopathology in Preadolescents

**DOI:** 10.1101/2021.10.14.464403

**Authors:** Xiang Xiao, Christopher Hammond, Betty Jo Salmeron, Hong Gu, Tianye Zhai, Hieu Nguyen, Hanbing Lu, Thomas J Ross, Yihong Yang

**Author notes:** To whom correspondence may be addressed: Yihong Yang, Ph.D. Neuroimaging Research Branch, National Institute on Drug Abuse, National Institutes of Health, Baltimore, Maryland, USA. **Email**.

## Abstract

**Background:** Cognitive dysfunction and high-order psychopathologic dimensions are two main classes of transdiagnostic factors related to psychiatric disorders. They may link to common or distinct core brain networks underlying developmental risk of psychiatric disorders.

**Method:** The current study is a longitudinal investigation with 11,875 youths aged 9-to 10-years-old at study onset, from the Adolescent Brain Cognitive Development study. A machine-learning approach based on canonical correlation analysis was used to identify latent dimensional associations of the resting-state functional connectome with multi-domain behavioral assessments of cognitive functions and psychopathological problems. For the latent rsFC factor showing a robust behavioral association, its ability to predict psychiatric disorders was assessed using two-year follow-up data and its genetic association was evaluated using twin data from the same cohort.

**Result:** A latent functional connectome pattern was identified that showed a strong and generalizable association with the multi-domain behavioral assessments (5-fold cross validation: ρ = 0.68~0.73, for the training set (N = 5096); ρ = 0.56 ~ 0.58, for the test set (N = 1476)). This functional connectome pattern was highly heritable (h^2^ = 74.42%, 95% CI: 56.76%-85.42%), exhibited a dose-response relationship with cumulative number of psychiatric disorders assessed concurrently and 2-years post-MRI-scan, and predicted the transition of diagnosis across disorders over the 2-year follow-up period.

**Conclusion:** These findings provide preliminary evidence for a transdiagnostic connectome-based measure that underlies individual differences in developing psychiatric disorders in early adolescence.

## Introduction

Over the past fifty years since the inception of the diagnostic and statistical manual (DSM) and the standardization of psychiatric nomenclature in the 1970s, psychiatry has focused on establishing diagnostic categories based upon clinical symptoms. The inability to identify biomarkers linked to these diagnostic categories has hindered progress in the field of psychiatry, which continues to struggle to reconcile nosologic problems related to comorbidity and specificity of current diagnostic categories. It is widely recognized that the diagnostic categories do not reflect unique underlying neuropathology. Emerging biological research points to shared genetic risk and overlapping structural and functional abnormalities across psychiatric disorders (1–4). This disconnection between current psychiatric nosology and biological findings highlights the need to examine neurobiological substrates and clinical symptoms that are shared across diagnoses.

Cognitive dysfunction is a common feature shared across diverse psychiatric disorders (5,6). To date, much of the research on cognitive dysfunction in psychiatric disorders has focused on identifying patterns of cognitive deficits within specific psychiatric disorder categories (e.g., schizophrenia) (7). Studies of cognition in population-based samples have identified an underlying, largely heritable, latent factor reflecting general cognition (g-factor) (8). A similar measure has been identified for psychopathology; the p-factor is a dimensional general psychopathology factor that cuts across disorder boundaries and is predictive of lifespan functional impairment and prospective psychopathology (9). While the relationships between cognition and psychopathology are complex and may be bidirectional (10), the fact that general cognition and psychopathology scores are commonly anticorrelated (9,10) and individually relate to function in overlapping brain regions, raises the question of whether they share neurobiological underpinnings.

Developmental factors play a large role in neural, cognitive, and behavioral manifestations of pediatric-onset psychiatric disorders. Three quarters of all psychiatric disorders emerge before the age of 21 years with 35% emerging before the age of 14 years (11). The presentation of psychiatric symptoms and disorders and brain structure and function change across the lifespan, exhibiting non-linear developmental trajectories, with marked shifts occurring during adolescence (12,13). In contrast, cognitive functions exhibit, in general, a more linear developmental trajectory, remaining relatively stable from childhood to young adulthood (14,15). If cognition dysfunction and psychopathology do share some neurobiological underpinnings, identifying these in children, many of whom will not yet have developed psychiatric diagnoses, may offer insight into neurobiological risk factors for the development of psychopathology.

Recent neuroimaging techniques, such as resting-state functional connectivity (rsFC) analysis, enable noninvasive investigation of the system-level organization of brain circuits via the temporal synchrony between brain regions (16). The functional connectome, the collective set of functional connectivity in the brain, can reliably discriminate one brain from another like a fingerprint (17), and is thought to underlie individual differences in cognitive and affective functions (17,18) and in expression or regulation of the psychopathological symptoms (19). Functional connectomes have been associated with a number of different psychiatric disorders (20) as well as with cognitive functions (17,21).

To better understand the co-aggregation of cognition and psychopathology and their shared and distinct neural substrates during development, in the present study, we sought to identify latent brain-behavior associations between the functional connectome and a broad set of behavioral assessments spanning the cognitive and psychopathological domains in preadolescent youth. We analyzed data from the Adolescent Brian Cognitive Development (ABCD) study (22,23), which includes brain imaging and comprehensive behavioral assessments from a large community-based sample of 9-10-year-old children in U.S., using canonical correlation analysis (CCA), a multi-view machine learning approach (24,25). CCA identifies latent components from two high-dimensional data sets that show maximal cross-set correlations (24,25). As a tool for brain-behavior association analyses, CCA discovers whole-brain connectivity patterns associated with a set of behavior assessments without *a priori* assumptions (26–28). This allows for the possibility of uncovering multiple ways in which brain measures may relates to behavioral constructs which can be identified as different modes. In this study, a rsFC pattern was identified that showed a significant and generalizable association with the behavioral assessments, positively correlating with cognitive functions while negatively correlating with psychopathological measures in a transdiagnostic manner. We repeated the CCA twice, once with only the cognitive measures and once with only the behavioral/emotional measures, and obtained essentially the same connectome variate, supporting the existence of a single underlying neurobiology supporting both cognitive and behavioral/emotional functioning. The rsFC pattern was highly heritable based upon analyses from a subgroup of twin-pair participants. The rsFC showed a dose-dependent relationship with cumulative number of psychiatric diagnoses assessed concurrently and two years post-fMRI scan, and predicted longitudinal transitions between healthy and diagnosed states over the 2-year study period, providing preliminary evidence of its potential usefulness in understanding risk for neuropsychiatric illnesses.

## Methods

### Participants

Neuroimaging data and behavioral assessments of 11,875 children aged 9-to 10-years were obtained from the ABCD study (22). This large and long-term ongoing project aimed to characterize psychological and neurobiological development from pre-adolescence to young adulthood. Participants and their families were recruited through school and community settings in 21 centers across the US, following locally and centrally approved Institutional Review Board procedures as detailed elsewhere (29).

From the 11,875 participants, 7,382 (3,714 females, aged 9.95 ± 0.62 y/o at baseline) met inclusion criterion, including 38 pairs of MZ twins (38 females, aged 10.30 ± 0.63 y/o at baseline) and 62 pairs of DZ twins (64 females, aged 10.16 ± 0.56 y/o at baseline). See Figure S1 for exclusion criteria, and Table S1 for demographic information of the included participants.

### Assessments of Multi-domain Behaviors

The behavioral dataset was comprised of multi-source data assessing cognitive functioning and psychopathology across multiple domains.

Participant’s cognitive functioning was assessed using performance data from a 15-domain neurocognitive test battery (40) and behavioral results from 3 neuroimaging tasks (41) (see Supplemental Table S2 for details). From the resultant data, 20 dimensional measures of cognition were obtained from each participant for the analysis.

Participant’s psychopathology at baseline was assessed using data from the parent-reported Child Behavioral Checklist (CBCL) and scale of mania, supplemented with self-report measures of impulsivity and psychosis risk (see Supplemental Table S3 for details). From the resultant data, 31 dimensional measures indexing different psychopathology-related constructs were obtained from each participant for the analysis. For all measures, baseline (7,382 participants) and 2-year-follow-up data (6,414 participants) were included into analyses.

### Clinical Diagnoses of Psychiatric Disorders

Participants were assessed at baseline and 2-year-follow-up for the presence of current and lifetime psychiatric diagnoses using the computerized-version of the Kiddie Schedule for Affective Disorders and Schizophrenia (K-SADS) for DSM-5 (KSADS-5), a psychometrically-validated semi-structured psychiatric interview (42,43).

### Resting-State Functional Connectome

In the ABCD Study, four 5-min resting-state functional image series and a T1-weighted structural image were collected for each participant with 3T scanners. Scanning parameters slightly differed across scanning sites and are detailed in Supplementary Materials.

Structural and functional MR images were preprocessed and housed in the ABCD-BIDS Community Collection (ABCC) from the Developmental Cognition and Neuroimaging (DCAN) Labs. The preprocessing pipeline included Human Connectome Project (HCP)’s minimal preprocessing pipeline (34) and the DCAN BOLD Processing (DBP) software (35). Processed data were obtained from collection 3165 provided by DCAN Labs. Descriptions of the pre-processing are detailed in Supplementary Materials.

The functional connectomes of individual brains were constructed from resting-state functional connectivity between 352 regions-of-interest (ROIs) across the brain. The ROIs were defined by the parcellation scheme from Gordon et al. 2016 (36), which included 333 cortical areas and 19 subcortical areas (Figure 1A). The ROIs were originally assigned into 12 functional communities (36) and further assembled into 7 large-scale networks according to the guideline by Uddin et al. (37). Functional connectivity was defined as the z-transformed Pearson’s correlation of the pre-processed BOLD time series between ROIs (Figure 1B). From the 352-by-352 matrix, the 61,776 upper-triangular values were used to describe the individual’s functional connectome.

**Figure 1.**
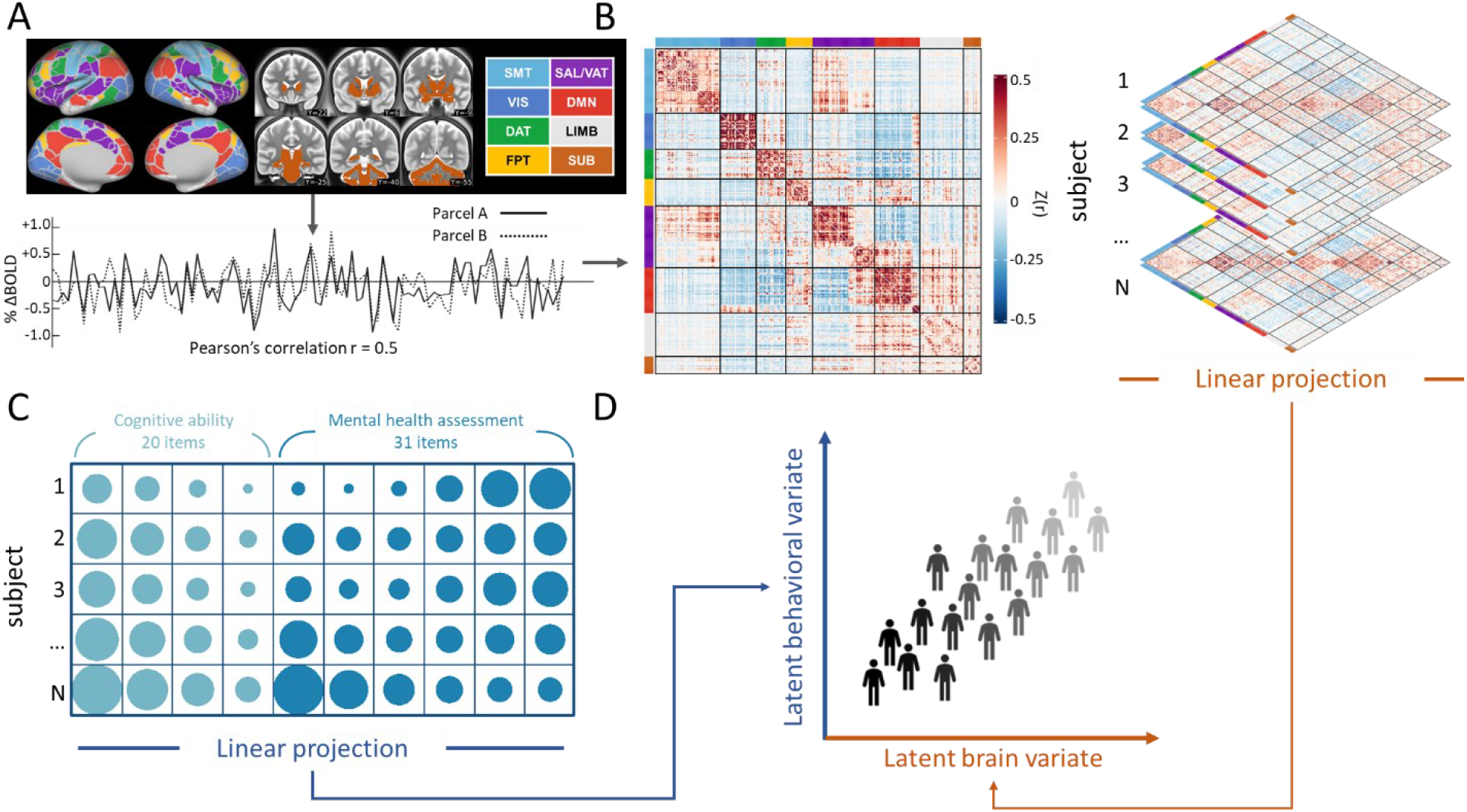
Schematic of the canonical correlation analysis (CCA). **(A) Construction of functional connectome.** Preprocessed time-series were extracted from 352 parcels. Then z-transformed time series correlations were calculated for each pair of parcels. Network abbreviations are listed in the SI. **(B) Connectome dataset.** Functional connectome data of 7382 participants (5906 as the training and 1476 as the test sets) were included as brain features**. (C) Behavioral-assessment dataset.** The behavioral assessments consisted of 20 scales for cognitive ability and 31 scales for mental-health related constructs. **(D) Population in the aligned latent space identified by CCA.** CCA identifies linear subspaces for the two datasets. The two datasets are ensured to be maximally correlated when projected to the latent spaces, and thus reveal latent brain-behavior associations.

### Covariates

To control for confounding factors, we considered two tiers of covariates that may affect the CCA analysis (38). Tier-1 included a set of technical variables related to MRI acquisition that may drive artificial connectome-behavior associations. These covariates were regressed out from both the connectome and behavior datasets. Tier-2 included a set of sociodemographic variables that may affect the connectome-behavior associations but with an unspecified causal mechanism. These covariates were used for a sensitivity analysis following the CCA. Descriptions of the covariates are detailed in Supplementary Materials.

### Discovery of Brain-Behavior Associated Dimensions with CCA

To identify associations between the functional connectome of the youth brain (Figure 1B) and multi-domain behavioral assessments (Figure 1C), we conducted a CCA (39) on the two datasets. To ensure the generalizability of the multi-variable result, the CCA was conducted in a hold-out framework (40); see Figure S2. The 7,382 participants were randomly split into a training set of 5906 subjects (80%) for discovery and a test set of 1476 subjects (20%) for validation. Within the training set, the CCA model was estimated. Both the connectome data and behavior data were controlled for the Tier-1 covariates and the residuals were transformed into a lower dimensional basis where data are exchangeable (23, 24). Then a reduction procedure was performed. A nested 5-fold hold out validation (80% vs. 20% for each fold) was performed to tune the hyperparameter, i.e., the number of principle components retained after dimension reduction (Figure S2 and Figure S3, see Supplementary Materials for more details).

CCA identifies orthogonal latent variates (Figure 1D) from the brain and behavior datasets, while ensuring a maximized correlation between the two variates paired by their orders (modes). To identify meaningful brain-behavior associations, we tested these modes for their statistical significance (41,42), generalizability (40), and redundancy index (26,43) via permutation tests. See Supplementary Materials for details about the statistical evaluation of the canonical variates.

### Association Between the Connectome Variate and Behavioral Assessments

To assess the identified association between the connectome variate and individual differences in cognitive performance and psychopathological measures, we first conducted a connectome-based prediction in a hold-out manner. Coefficients of the first CCA mode estimated from the training set, produced by multiplying the weighting matrices given by PCA and CCA, were applied to the test set rsFC data. Pearson’s correlations between the predicted connectome variate score to each behavioral assessment were calculated. Then for a prospective prediction, the connectome variate scores were used to predict the behavioral assessments in the 2-year-follow-up data. P values were corrected for multiple comparisons using the Benjamini-Hochberg false discovery rate (FDR) method (44).

### Association Between the Connectome Variate, Clinical Diagnoses, and Status Transitions

With the goal of determining the clinical utility of the connectome variate in predicting psychiatric disorders, we characterized associations between the connectome variate at baseline and clinical diagnoses at baseline and 2-year-follow-up. To examine disorder-specific relationships: the 7382 participants were grouped by their K-DSADS diagnoses, and the connectome variate score of each diseased group was compared with the group with no current psychiatric diagnosis. Three analyses were conducted to examine transdiagnostic relationships: First, participants were grouped by 0, 1, 2, or ≥3 comorbid diagnoses. ANOVAs with post-hoc Games-Howell comparison procedure were used to compare differences between groups. Second, the risk of at least one diagnosis was compared between participants with a connectome variate score above MEAN+1SD vs. under MEAN-1SD in the whole cohort. Lastly, using data on current psychiatric diagnoses at baseline and 2-year-follow-up, we stratify participants into four ‘transition’ groups of Healthy-persistent, Disorder-persistent, Disorder-remitted, Disorder-new-onset (see Figure 8A and supplemental methods for more details). Then we used ANOVAs to compare connectome variate scores across transition groups.

**Figure 2.**
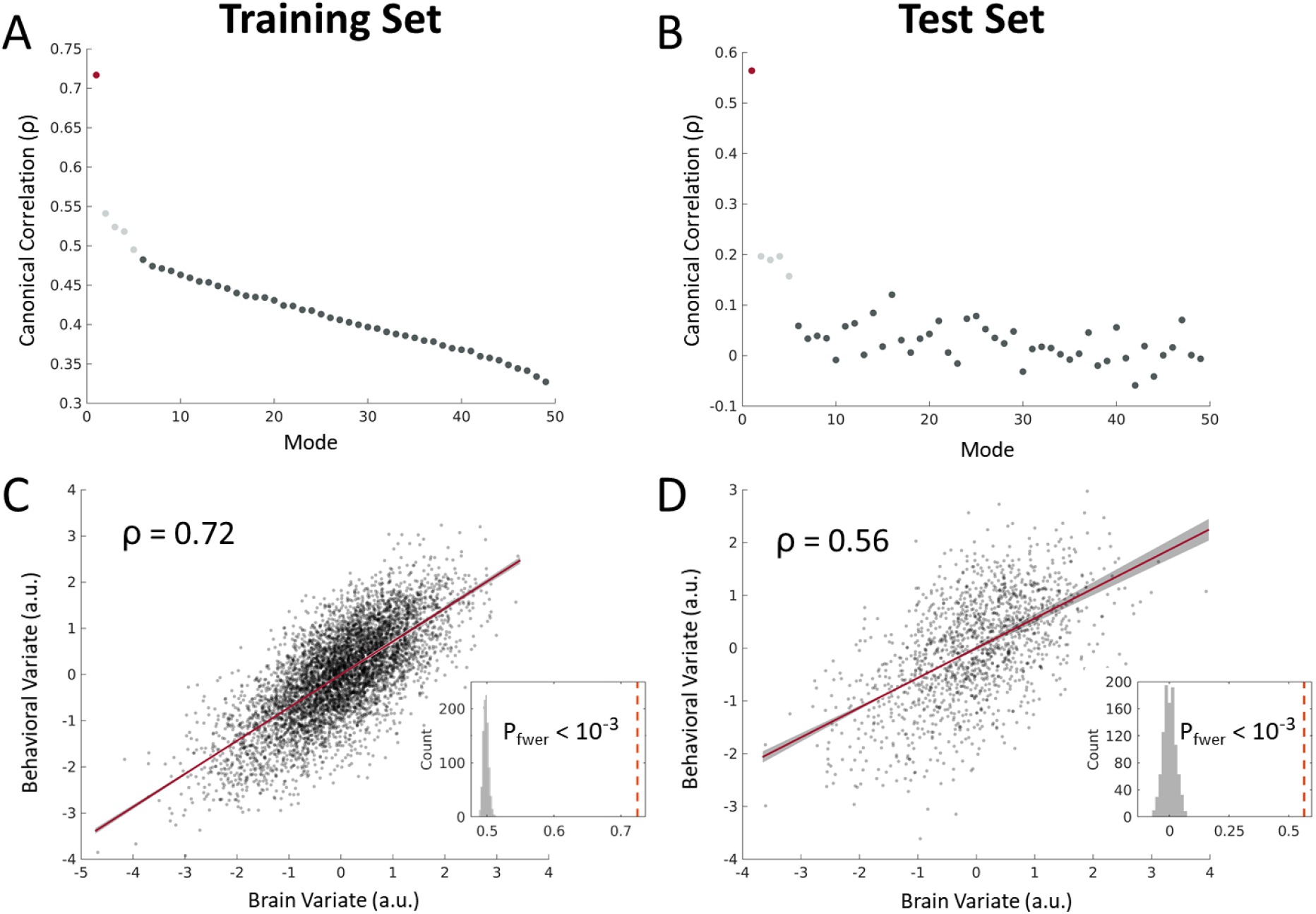
Canonical correlations (CCs) between the brain and behavioral datasets. **(A)** CCs of the 49 CCA modes from the training set. In the training set, top five (red and light gray points) modes show significant correlations in comparison to the null distribution. **(B)** CCs of the 49 modes from the test set. When generalizing to the test set, the top five CCA mode shows significant CCs in comparison to the null distributions five (red and light gray points). **(C) and (D)** Scatter plots of the associated brain and behavior variate scores in mode 1, for training and test sets respectively. Inset plots represent permutation test results conducted for CC in the training and test sets

**Figure 3.**
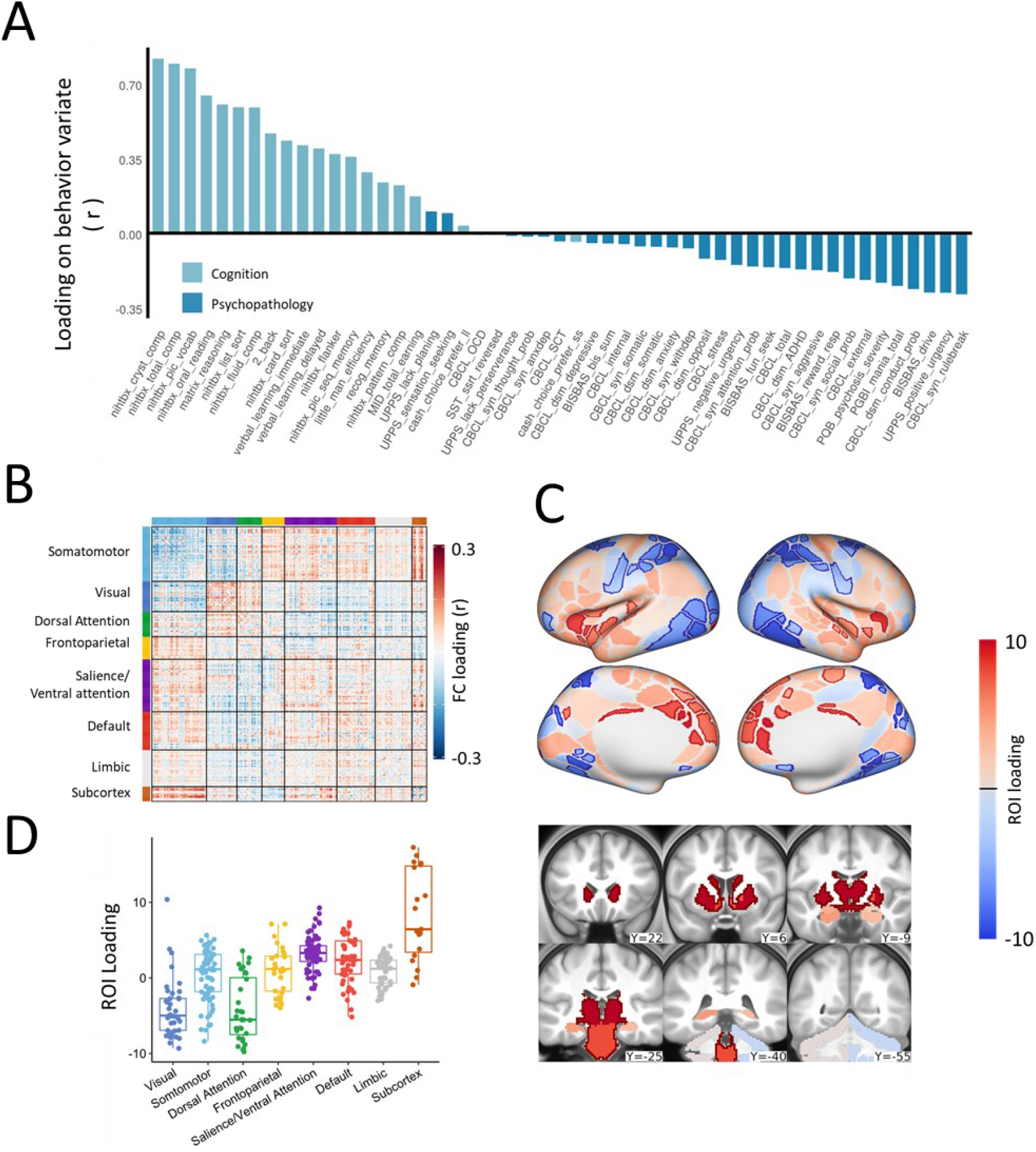
Brain and behavioral loadings of the first CCA mode. **(A) Loading of each behavioral assessment on the behavioral variate.** Abbreviations are listed in Tables S2-S3 **(B) Loading of rsFC on the brain connectome variate. (C) Map of ROI loading**. Highlighted borders indicate ROIs with significant positive (dark red) or significant negative (dark blue) loadings (p < 0.01, FDR-corrected for multiple comparisons). **(D) Distribution of ROI loadings across large-scale functional brain networks.** The 352 parcels were grouped into 7 cortical networks and the subcortex (*47*).

**Figure 4.**
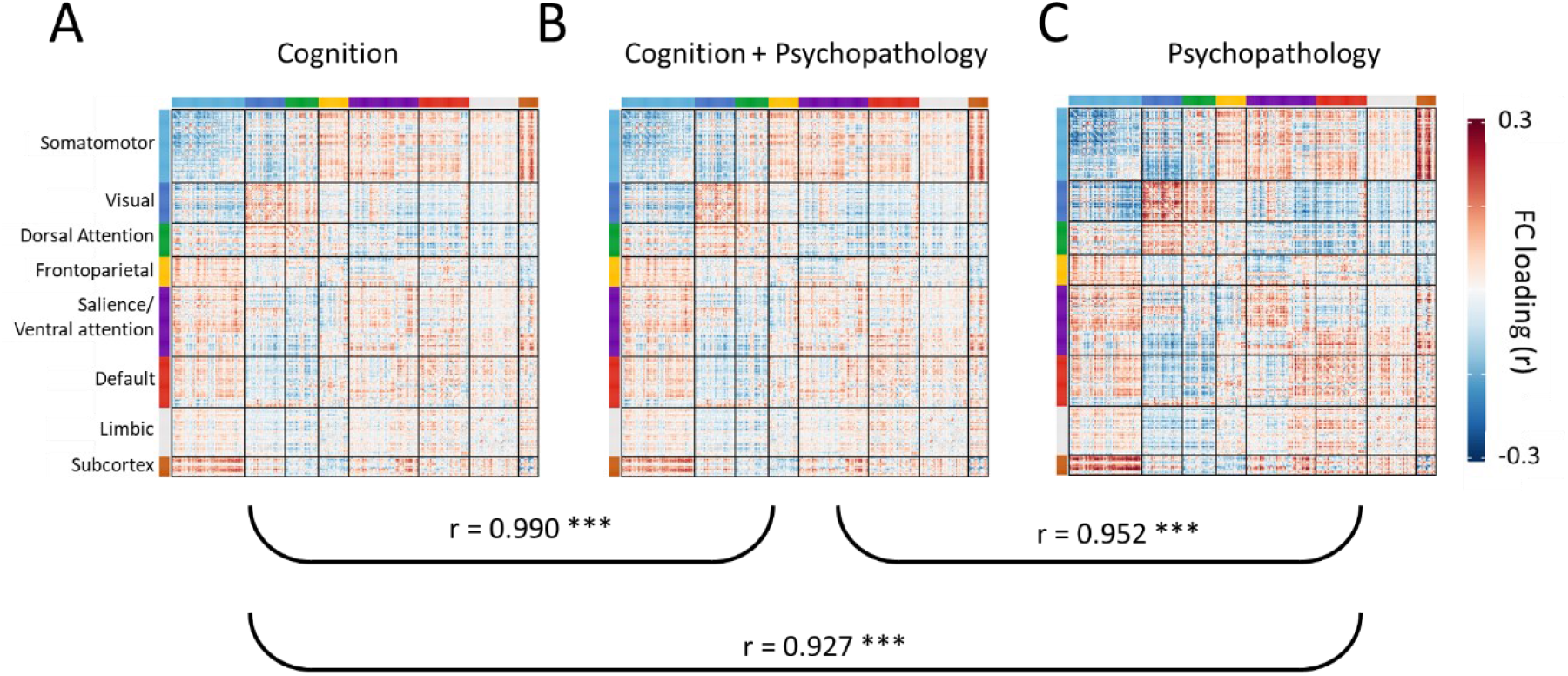
Cross-domain consistent connectome variate of the first CCA mode. Loading maps of the first CCA mode obtained by associating the connectome data set to the **(A)** Cognition only **(B)** Combined cognition and psychopathology and **(C)** Psychopathology only behavioral set show high consistency (spatial Pearson’s correlation) between them. Loading maps are averaged across the cross-validation folds. The resulting p-values are corrected for multiple comparison using the FDR procedure, *** FDR-adjusted p < 0.001.

**Figure 5.**
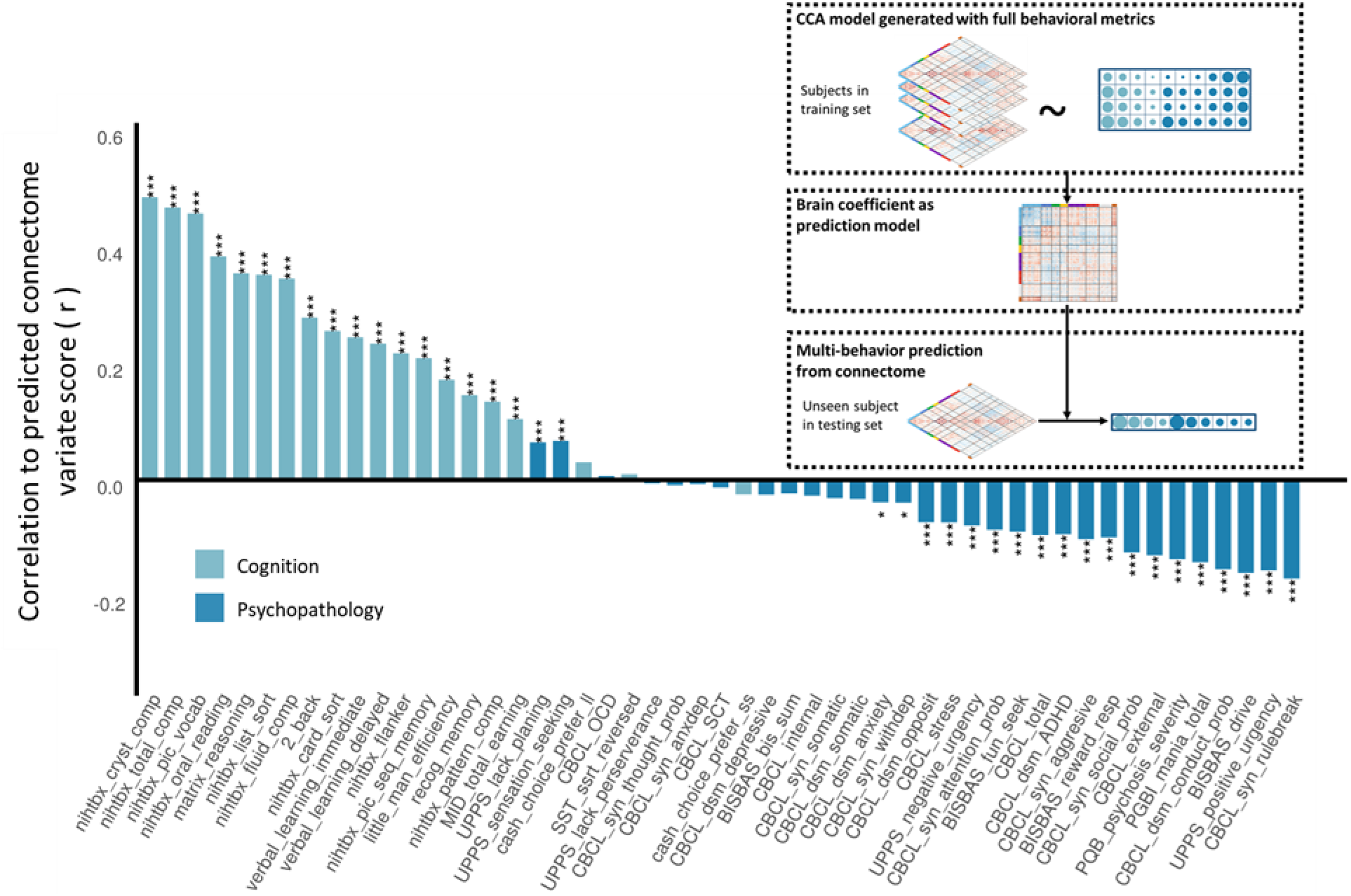
Behavioral assessment prediction from the connectome variate score. The bar plot shows the performance of using the CCA estimated connectome variate to predict behavior assessment scores (ordered by loading) in an unseen population. Inset shows the cross-population prediction procedure. The predictive ability of behavioral assessments was assessed with Pearson’s correlation. FDR-adjusted p-values: * *p* < 0.05, ** *p* < 0.01, *** *p* < 0.001

**Figure 6.**
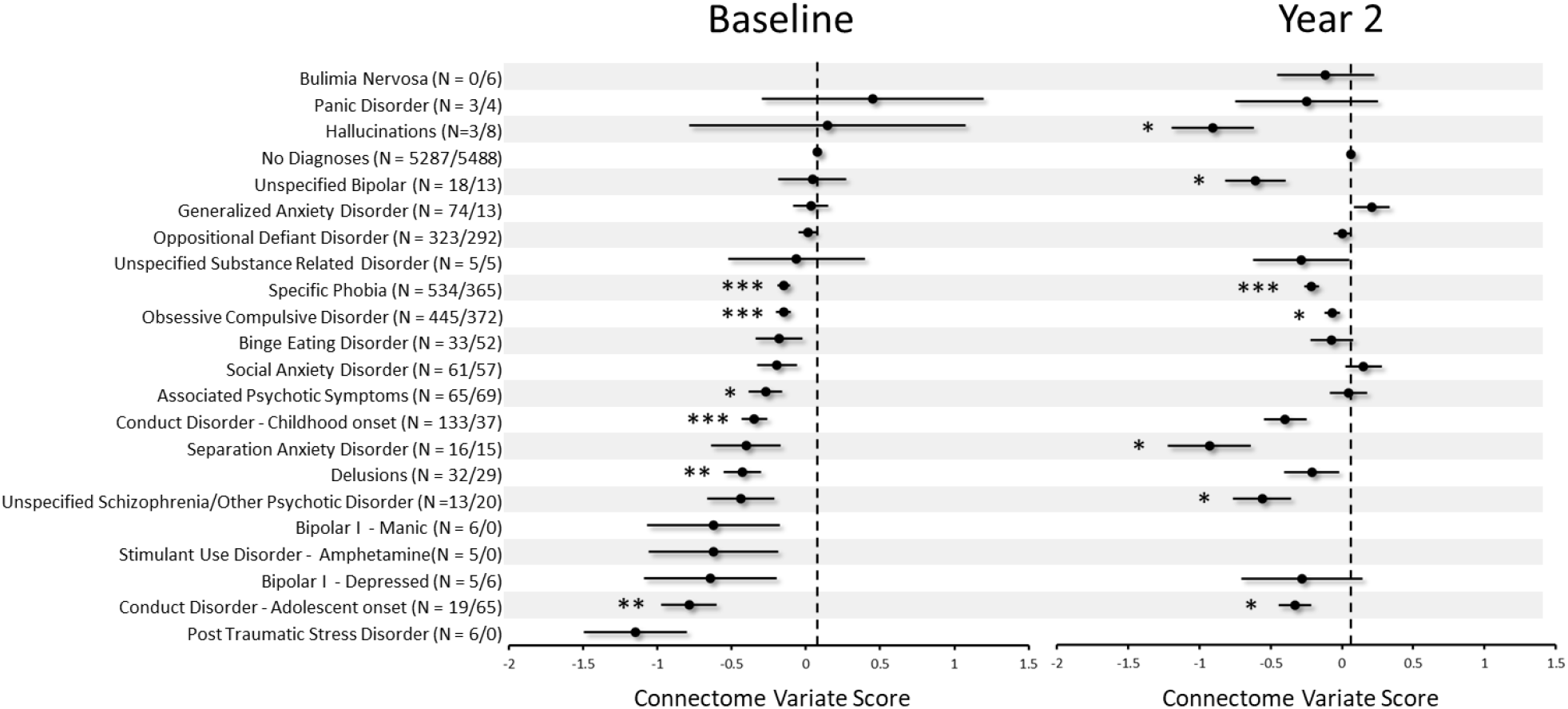
Relationships between connectome variate scores and occurrence of psychiatric disorders in the entire sample, at baseline and the 2-year follow up. Error bars indicate standard error. Welch’s *t*-tests were used to compare differences between each diagnostic subgroup to the control (i.e., no diagnoses) group. FDR-adjusted p-values: * *p* < 0.05, ** *p* < 0.01, *** *p* < 0.001

**Figure 7.**
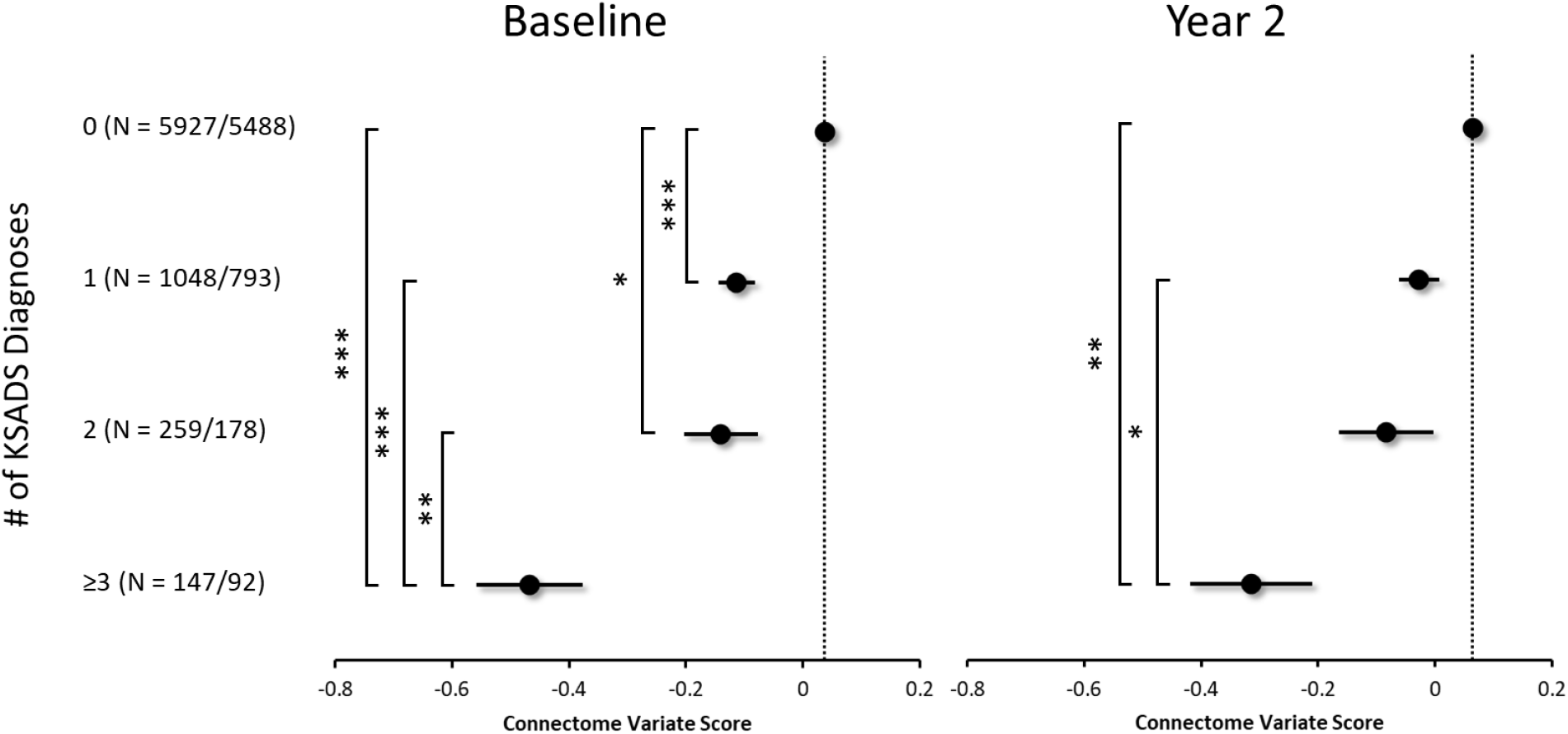
Relationships between connectome variate scores and the number of simultaneous presenting psychiatric disorders. Group-wise comparison of mean connectome variate scores in preadolescents with the cumulative number of comorbid disorders at baseline and 2 years after the MRI scan. Dots and lines in the plot indicate mean and standard error of the connectome variate score in each group of comorbid KSADS diagnosis. Pairwise group differences were post-hoc compared using the Games-Howell procedure. * Adjusted *p* < 0.05, ** Adjusted *p* < 0.01, *** Adjusted *p* < 0.001

**Figure 8.**
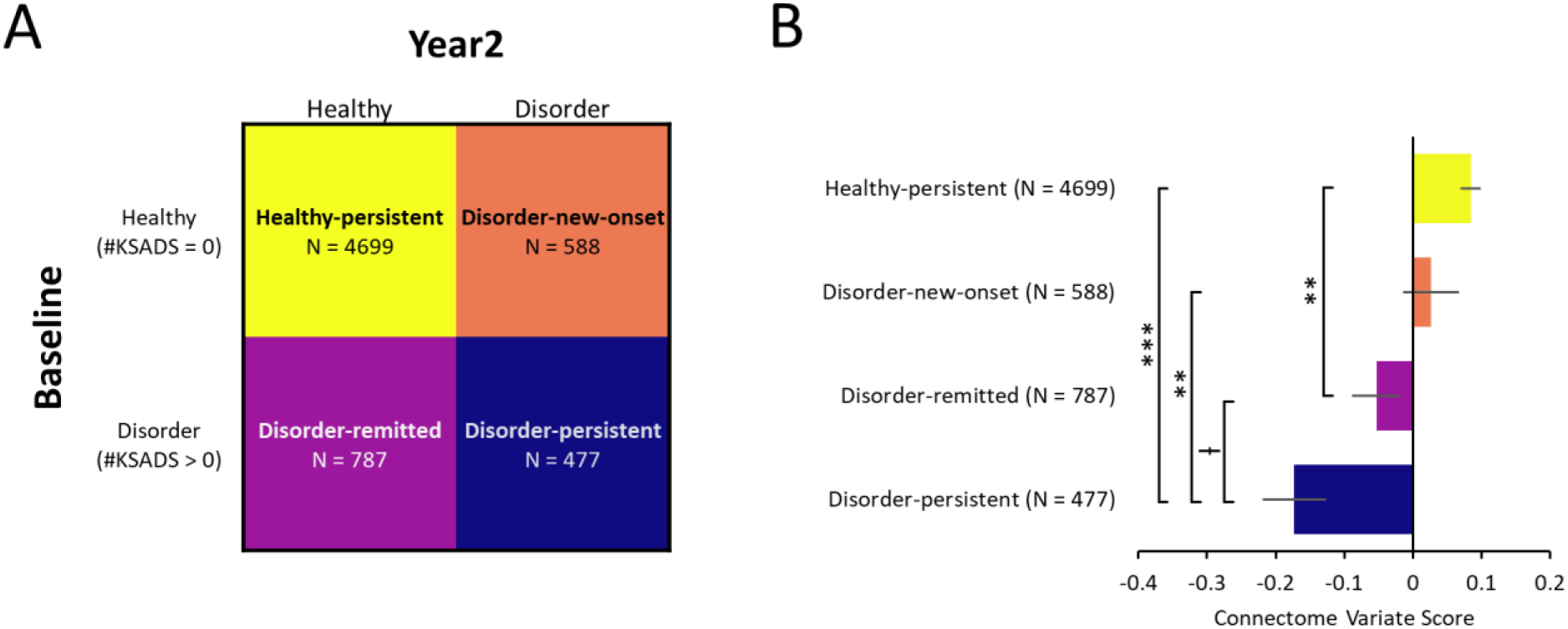
Relationships between connectome variate scores and the longitudinal transition of KSADS diagnosis status. **A) Groups defined according to the longitudinal transition of KSADS diagnosis status. B) Comparison between the groups.** Bars and error bars indicate mean and standard error of the connectome variate score in each group of comorbid KSADS diagnosis. Pair-wise group differences were post-hoc compared using the Tukey’s procedure. ł unadjusted *p* <0.05, * Adjusted *p* < 0.05, ** Adjusted *p* < 0.01, *** Adjusted *p* < 0.001

### Heritability of the Connectome Variate

Twin data from the 7,832 participants were used to estimate the heritability of the identified connectome variate. Linear structural equation model with a classic twin study design (CTS) (46) was used. The total phenotypical variance, P, was attributed into a sum of genetic and environmental factors (P = A + C + E): additive genetic influences (A), environmental influence shared by family members (common environmental variance, C) and unique environmental influence (E). Goodness of fit (GOF) was tested for the full ACE model and its sub-models (models restricting path(s) to be zero). Akaike’s information criterion was used to determine the best model considering both the GOF and the model parsimony. Significance of each path was assessed by comparing the fit of the restricted model to the full model. A chi-square test was used for statistical assessment. Heritability (h^2^) was estimated as the ratio of the genetics related variance to the total phenotypical variance.

## Results

### Latent Dimensions Linking the Brain Connectome with Cognition and Psychopathology

CCA identified 49 connectome-behavior association modes (pairs of latent connectome and behavioral variates showing correlation). The first 5 modes exhibit significant connectome-behavior associations (Table S4, Figure 2A and B), both in the training and test set, compared to the null distribution generated by permutation (see Supplementary Materials).

The first mode (ρ = 0.72, family-wise error rate [fwer] adjusted *p*_fwer_ < 10^-4^ in the training set; and ρ = 0.56, permutation *p*_fwer_ < 10^-4^ in the test set; shown in Figure 2C and Figure 2D, respectively), uniquely accounted for a large proportion of redundant variance (43) between the brain and behavior datasets (Table S4). Further, its associated connectome variate, described below, manifests a striking cross-domain consistency. Thus, we focus on the first mode in the following results (See Figure S4 and S5 for other significant modes).

Figure 3A shows the loading of each behavioral assessment on the latent behavioral variate of the first CCA mode. Nearly all assessments related to cognitive ability show positive loadings on the latent behavior variate, whereas most of the psychopathology-related constructs show negative loadings. Furthermore, differences are observed across externalizing and internalizing domains of psychopathology. Measures indexing the externalizing ‘spectrum’ problems generally exhibit higher loadings (e.g., *r* = −0.27 for CBCL Rule Breaking and *r* = −0.25 for CBCL Conduct Problem), compared to the internalizing ‘spectrum’ problems (e.g., *r* = −0.008 for CBCL Anxious-Depression and *r* = −0.06 for CBCL Withdrawn-Depression).

Figure 3B shows the rsFC loading, defined as correlation between rsFC and the connectome variate of the first CCA mode for each ROI pair. A positive loading indicates that the rsFC of the ROI pair contributed positively to the association in which higher brain connectome variate was correlated with better performance in cognitive tasks and lower severity of parent-reported psychopathological symptoms. A measurement of ROI loading is defined by summing the rsFC loadings of that ROI (i.e., each row in the loading matrix in Figure 3B), indicating the overall contribution of the rsFC associated with the given ROI to the identified brain connectome variate. This measure reveals that the rsFC loadings distribute unequally (Gini index = 0.185 and 0.181 for positive and negative loadings, permutation test *p* < 0.0001 in both cases) across the whole-brain ROIs (see Figure S7). ROIs formed two clusters, each predominantly significant positive or negative rsFC loadings (Figure 3C), compared to the null distribution of ROI loadings generated by permutating the loading matrix of Figure 3B.

ROIs with significant positive loadings include cortical areas of the anterior insula/frontal operculum (aI/fO), dorsal anterior cingulate cortex (dACC), ventral lateral prefrontal cortex (vlPFC), dorsal medial prefrontal cortex (dmPFC), superior temporal gyrus (STG), as well as subcortical regions of the caudate, putamen, accumbens, thalamus and brain stem. ROIs with significant negative loadings encompass cortical regions of the intra-parietal sulcus and superior parietal lobule (IPS/SPL), frontal eye fields (FEF), as well as the pre-motor area (PMA), and associative visual areas in the occipital lobe (Figure 3C).

Furthermore, the loading of the ROIs significantly differed by their affiliation to large-scale functional networks (Figure 3D; one-way ANOVA, *F*(7,344) = 42.58, *p* < 0.001). Specifically, negative loadings appeared mainly in the visual (VIS) and the dorsal attention network (DAN), while the largest positive loadings were mainly presented in the salience/ventral attention network (SAL/VAN), default mode network (DMN), and subcortical network (SBC).

Finally, the results for the first CCA mode replicated when sociodemographic covariates were controlled for (Figure S8), with generalized canonical correlations of *ρ* = 0.48~0.58, and similar connectome variate loadings (*r* = 0.86~0.99) and behavioral loadings (*r* = 0.98~0.99) comparing with the main result.

### Cross-Domain Consistency of the Mode-1 Connectome Variate

The transdiagnostic association revealed by the first CCA mode implies that deficits of cognitive functions and behavioral/emotional problems may share a common set of neural processes. To exclude the alternative that such an association may be driven by only one domain, we conducted the same CCA procedure described above twice, but with behavioral assessments from only a single domain, cognition or behavioral/emotional, at a time. In both cases, the first mode survived from both tests of within-sample significance and out-of-sample generalizability (Figure S7). Although the canonical correlation values differed between the CCA models estimated from cognition, behavioral/emotional (Figure S7) or their combination (Figure 2), the connectome variates showed very high consistency among the three cases (combined vs. cognition only, *r* = 0.990; combined vs. behavioral/emotional only, *r* =0.952; Figure 4). Note that cross-domain consistency with high correlation (i.e., |*r*| > 0.9) was found to be specific to the first mode (see Table S5 for other significant modes).

### The Connectome Variate Predicts Multi-Domain Behavioral Assessments

We next projected the mode-1 connectome variate estimated from the training set onto the independent test dataset, using rsFC to predict a wide range of univariate behavior assessments across the cognition and psychopathology domains.

Specifically, all cognitive domain assessments except the cash choice task and stop-signal reaction time were significantly predicted (FDR-adjusted *p* < 0.05). The psychopathology related constructs significantly predicted (FDR-adjusted *p* < 0.05) included measures of externalizing problems: e.g., CBCL Rule-breaking Behavior, CBCL Aggressive Behavior, and CBCL externalizing subscale; internalizing problems: e.g., CBCL Somatic Problems, CBCL Withdrawn/Depressed; and other problems: e.g., CBCL Social Problems, CBCL Oppositional Defiant Problems, CBCL ADHD, CBCL Stress, UPPS-P Positive Urgency, UPPS-P Negative Urgency, P-GBI Mania, and PQ-B Psychosis Severity. While the analysis showed that the connectome variate predicted psychopathology across domains, measures in the externalizing domain tended to have larger effect sizes than internalizing, and the CBCL externalizing subscale is significantly predicted while the CBCL internalizing subscale is not. See Figure 4 for all assessments.

Critically, these predictions of behaviors held for the first mode from CCA models trained on the cognition behavioral domain or psychopathology-related constructs separately (Figures S9 and S10).

Finally, the connectome variate score at baseline prospectively predicted the behavioral assessments at the 2-year follow up (Table S6).

### The Connectome Variate Predicts Clinical Diagnoses and Status Transitions

For analyses characterizing relationships between the connectome variate and clinical diagnoses, transdiagnostic and diagnosis-specific associations were found, both concurrently and at 2-year-follow-up.

In diagnosis-specific analyses comparing participants stratified by specific psychiatric diagnoses, a pattern emerged whereby lower connectome variate scores were associated with the morbidity of various current psychiatric disorders at baseline and at the 2-year follow up (Figure 6).

Participants were stratified into groups (0, 1, 2 and ⩾3) based on the cumulative number of current psychiatric disorders at each timepoint. Lower connectome variate scores were associated with more comorbid psychiatric disorders both at baseline (one-way ANOVA, *F*(3,7377) = 19.86, *p* <0.001) and at the 2-year follow-up (one-way ANOVA, *F*(3, 6547) = 7.25, *p* < 0.001); see Figure 7.

In our second transdiagnostic analysis (see Table S7), compared to those with high connectome variate scores (⩾1 SD above MEAN), participants with low connectome variate scores (⩽1 SD below MEAN) had a higher risk of having more than one psychiatric disorder at baseline (Risk Ratio = 1.76, 95% CI:1.36-2.29, *p* < 0.001) and at the 2-year follow up (Risk Ratio = 1.33, CI:1.09-1.62, *p* =0.004).

Results from our longitudinal transition analysis, which compared participants stratified based upon their pattern of change in current psychiatric diagnoses between diagnosed and healthy states over time (see Figure 8A), showed a significant association between the connectome variate score and transition group membership (One-way ANOVA *F*(3,6547) = 12.99, *p* < 0.0001; see Figure 8B).

### Heritability of the Connectome Canonical Variate

Intrapair correlations for the connectome variate score are *r*_MZ_ = 0.71 for MZ twins and *r*_DZ_ = 0.39 for DZ twins (Table 1). The AE model best fit the twin data, with a heritability of the connectome variate of h^2^ = 74.42% (95% CI: 56.76%-85.42%).

**Table 1.** Intrapair twin correlations, influence of genetic/environmental factors, heritability of connectome variate.

| Twin Correlations |  | Variance |  | Heritability |
| --- | --- | --- | --- | --- |
| $r_{MZ}$ (N) | $r_{DZ}$ (N) | $a^2$ (95% CI) | $e^2$ (95% CI) | $h^2$ (95% CI) |
| 0.71 (76) | 0.39 (126) | 0.60 (0.42 -0.82) | 0.20 (0.13 - 0.32) | 0.74 (0.56-0.86) |

## Discussion

Applying CCA to a multidimensional dataset that included comprehensive measurements of rsFC, cognition, and psychopathology from a large community-based sample of U.S. preadolescents, we identified a single connectome-based latent brain variate that covaried with a broad number of cognitive and psychopathological measurements. The connectome variate had rsFC loadings distributed in attention and cognitive control networks. The connectome variate was positively correlated with cognitive performance across domains and negatively correlated with behavioral/emotional measures across dimensions and diagnoses. It is notable that the connectome variate could be robustly identified with either of the domains. The brain variate predicted a range of behavioral measures and clinical diagnoses including dose-dependently predicting the cumulative number psychiatric disorders at baseline and at the 2-year follow up. Finally, the connectome variate was highly heritable, suggesting that it may reflect intrinsic differences with genetic underpinnings. Together, these findings provide preliminary evidence for a connectome-based biomarker that indexes individual differences in cognitive dysfunction and transdiagnostic risk of developing psychopathology, with higher variate scores reflecting resilience and lower scores reflecting vulnerability.

Identification of psychiatric biomarkers is essential for both understanding the ontology of psychiatric disorders and for developing precise treatments for them. Traditional efforts to identify psychiatric biomarkers have relied predominantly on case-control designs and focused on neural alterations within a single disease. However, few clinically-useful disorder-specific brain-behavior associations have been discovered using these approaches. This has led to a shift in approach towards identifying core brain-behavior associations that cross traditional diagnostic boundaries similar to the approach promoted by the RDoC initiative (48,49). Neuroimaging studies using a transdiagnostic framework have demonstrated consistent findings showing spatially overlapping neural deficits, including lower gray matter volume (3) and impaired activation during cognitive tasks (50), across diagnostic categories. Consistent with these findings, results of the current study identified differences in rsFC across large-scale brain networks, suggestive of maladaptive connectomes, that are associated with a broad spectrum of psychopathology and cognitive processes, and may represent a transdiagnostic neural marker for psychiatric disorders. However, while the RDoC approach seeks associations with specific cognitive and affective functions across diagnoses, we found a single variate that related, albeit to differing degrees, to most aspects of both cognition and psychopathology, irrespective of diagnosis. In addition, given the ABCD sample, our findings extend the developmental window for observing transdiagnostic neural deficits from adulthood down to middle childhood, a critical period for the onset of mental disorders. As such, our connectome-based variate may provide unique predictive or prognostic value related to the emergence and progression of psychiatric disorders from preadolescence to young adulthood, providing a roadmap for developing brain-based treatment targets.

It is notable that we derived essentially the same brain variate when using the cognitive or psychopathologic features in isolation and when using them in combination, suggesting that the two behavioral domains share essential neural processes. Our findings parallel prior epidemiologic research showing that general cognition and psychopathology factors are anticorrelated (51,52). Our results extend these findings by showing that measures of cognition and psychopathology that have traditionally been viewed as separate constructs, load on the positive and negative ends of the same dimensional factor (rather than orthogonal ones) and covary with the same connectome variate. The non-specific association between our connectome-based brain variate and wide array of psychopathology measures and psychiatric diagnoses, suggests that this brain signal indexes transdiagnostic rather than disorder-specific vulnerability in preadolescents. That the brain variate exhibits a dose-response relationship with cumulative number of current psychiatric disorders suggests potential clinical applicability as a diagnostic-severity staging neural index, and warrants further exploration. While our findings require cautious interpretation until out of sample validation and more prospective testing is performed, they provide early support for a latent connectome-based brain variate being a candidate biomarker indexing individual differences in risk for transdiagnostic psychopathology in preadolescents.

In addition to its efficiency in explaining individual variance shared by multiple behaviors, our brain-based variate mirrors general behavior-derived factors like g-(53–55) and p-(56–58) in their heritability. Note that the general behavioral factors are genetically associated to each other (55), suggesting that the two traits may tap a shared neural process rooted in gene. In the current twin data, the connectome variate exhibited high heritability, indicating the special connectome pattern may act in the gene-environment correlation process as a mediator (59), facilitating one’s ability to cope with negative environment during the development.

Our rsFC loadings identified diverse cortical and subcortical brain regions including the aIns/fO, dlPFC, vmPFC, STG, IPS/SPL, FEF, PMA, striatum, thalamus and brain stem that contributed significant variance to the connectome-based brain variate. Notably, many of these cortical regions map onto frontoparietal brain systems which include the VAN/SAL and DAN, which are implicated in attention-based cognitive control (60–64). As such, proper functioning of these networks may be required to support healthy cognitive performance and protect against the development of psychiatric disorders (19), and alterations of rsFC in these ROIs may result in both cognitive dysfunction and the emergence of psychopathology. We found that significant ROIs form two clusters according to the direction of their loadings. Specifically, negative loadings were mainly located in the DAN while positive ones in the VAN/SAL, which may be aligned with the distinct functions of these two subsystems in supporting attention-based cognitive control (61,65). Subcortical regions including the striatum, thalamus, and brain stem also had significant positive loadings. This may indicate the critical role of these regions in supporting cognition by relaying, shifting, and sustaining functional information from their cortical counterparts (66).

This study has some noteworthy limitations. As the study is a secondary analysis of ABCD data, we were reliant on the assessments, data collection procedures, imaging parameters, and acquisition and harmonization choices made by the primary study team. While broad psychopathology indices were collected, they were mostly restricted to parent-reported measures at this timepoint, which is limiting and should be cross-validated for accuracy with multi-informant assessments from subsequent data waves. While our report emphasizes transdiagnostic psychopathology, it is important to note that most study participants scored within normal range on both psychopathological and cognitive assessments and did not meet criteria for psychiatric diagnoses at either timepoint. Our data-driven analytic approach also has limitations. CCA was designed to detect linear associations between the two multi-dimensional datasets of functional connectome and behavioral assessments. Non-linear models such as kernel CCA (24) might be valuable for identifying other linkages between functional connectome and the multi-domain behaviors, which would be important for finding biotypes within specific clinical categories or cognitive domains.

In conclusion, by applying a machine-learning CCA approach to multimodal brain and behavioral data from a large community sample of preadolescents who participated in the ABCD study, we identified a connectome-based variate that indexes individual differences in cognitive functioning and transdiagnostic psychopathology risk in this population. Our findings provide early evidence that system-level brain mechanisms may underly a common pathway conveying risk versus resilience in relation to development of psychiatric disorders.

## Acknowledgments

This research was supported by the Intramural Research Program of the National Institute on Drug Abuse, National Institutes of Health.

Data used in the preparation of this article were obtained from the Adolescent Brain Cognitive Development (ABCD) Study (https://abcdstudy.org), held in the NIMH Data Archive (NDA). This is a multisite, longitudinal study designed to recruit more than 10,000 children aged 9-10 years and follow them over 10 years into early adulthood. The ABCD Study is supported by the National Institutes of Health and additional federal partners under award numbers U01DA041048, U01DA050989, U01DA051016, U01DA041022, U01DA051018, U01DA051037, U01DA050987, U01DA041174, U01DA041106, U01DA041117, U01DA041028, U01DA041134, U01DA050988, U01DA051039, U01DA041156, U01DA041025, U01DA041120, U01DA051038, U01DA041148, U01DA041093, U01DA041089, U24DA041123, U24DA041147. A full list of supporters is available at https://abcdstudy.org/federal-partners.html. A listing of participating sites and a complete listing of the study investigators can be found at https://abcdstudy.org/consortium_members/. ABCD consortium investigators designed and implemented the study and/or provided data but did not necessarily participate in analysis or writing of this report. This manuscript reflects the views of the authors and may not reflect the opinions or views of the NIH or ABCD consortium investigators.

The ABCD data repository grows and changes over time. The ABCD data used in this report came from NDA Study 901. DOIs can be found at DOI 10.15154/1519007.

This work utilized the computational resources of the NIH HPC Biowulf cluster (http://hpc.nih.gov).

