## Supplementary Materials for "Brain Functional Connectome Defines a Transdiagnostic Dimension Shared by Cognitive Dysfunction and Psychopathology in Preadolescents"

*Supplementary methods*

**Criteria for Participants Exclusion**

As shown in Figure S1, participants were excluded due to missing scanning information, neurological condition, unavailability of the preprocessed MRI data, failure to pass Freesurfer quality control, or insufficient fMRI time points (<6 min) after motion censoring. As a result, data of 7382 participants (3714 females, aged 9.95 ± 0.62 y/o) were included in the final analysis. Among the 7382 participants, 38 pairs of MZ twins (38 females, aged 10.30 ± 0.63 y/o at baseline) and 62 pairs of DZ twins (64 females, aged 10.16 ± 0.56 y/o at baseline) are included.

Demographic information of the included participant is summarized in Table S1.

**Assessments of Cognitive Function**

The cognitive ability of the participants was quantified with 20 scores derived from their performance on 14 available neuro-cognitive tests (1) and 3 neuroimaging tasks (2).

The NIH Toolbox cognition battery includes seven tasks measuring multiple aspects of cognition, which generates the following scores: Picture Vocabulary Test Score (nihtbx_pic_vocab); Flanker Inhibitory Control and Attention Test Score (nihtbx_flanker); List Sorting Working Memory Score (nihtbx_list_sort); Dimensional Change Card Sort Test Score (nihtbx_card_sort); Pattern Comparison Processing Speed Test Score (nihtbx_pattern_comp); Picture Sequence Memory Test Score (nihtbx_pic_seq_memory); Oral Reading Recognition Test Score (nihtbx_oral_reading). From the direct scores of these tests, the NIH Toolbox further derives 3 higher-order composited scores: Cognition Fluid Composite Score (nihtbx_fluidcomp); Crystallized Composite Score (nihtbx_cryst_comp); and Cognition Total Composite Score (nihtbx_total_comp). For the above measures, uncorrected raw scores were used due to the undergoing revision of the NIH Toolbox (1,3).

Measurements and abbreviations of cognitive functions are listed in Table S2.

**Assessments of Psychopathology Related Constructs**

In the current study, 31 dimensional scores of psychopathology-related constructs in youth derived from baseline assessments were used to estimate the CCA based brain-behavior association. The measurements include the parent/caregiver reported Child Behavioral Checklist (CBCL), the scale of mania, and the youth reported scales of impulsivity and psychosis risk.

The parent/caregiver-reported CBCL (4) was used to assess the emotional and behavioral problems of the participants. It includes 119 problem items rated with 3-level scales: 0 = Not True, 1 = Somewhat or Sometimes True, and 2 = Very True or Often True. From the rating on these items, the CBCL further derives composite scales including: 8 syndrome scales of Anxious/Depressive symptoms (CBCL_syn_anxdep), Withdrawn/Depressed symptoms (CBCL_syn_withdep), Somatic Complaints (CBCL_syn_somatic), Social Problems (CBCL_syn_social_prob), Thought Problems (CBCL_syn_thought_prob), Attention Problems (CBCL_syn_attention_prob), Rule-Breaking Behaviors (CBCL_syn_rulebreak) and Aggressive Behaviors (CBCL_syn_aggresive); 3 higher-order scales of Internalizing Problems (CBCL_internal), Externalizing Problems (CBCL_external) and Total Problems (CBCL_total); 6 DSM-oriented scales of Depressive Problems (CBCL_dsm_depressive), Anxiety Problems (CBCL_dsm_anxiety), Somatic Problems (CBCL_dsm_somatic), Attention Deficits (CBCL_dsm_ADHD), Oppositional Defiant Problems

(CBCL_dsm_opposit), Conduct Problems (CBCL_dsm_conduct_prob) and 3-scale scores of Sluggish Cognitive Tempo (CBCL_SCT), Obsessive-Compulsive Problems (CBCL_OCD) and Stress Problems (CBCL_stress). For each of the 20 scales, raw scores were used in the current analysis.

The parent/caregiver-reported ten-item Mania Scale (5) derived from the 73-item Parent General Behavior Inventory (P-GBI) for Children and Adolescents (6) was used to assess dimensional mania symptoms of the youth. The sum score (PGBI_mania_total) was included in the current analysis.

A youth-report version of UPPS-P (Urgency, Perseverance, Premeditation and Sensation seeking) scale was used to assess the dimensional impulsivity of children (7,8). Five dimensional scales of Lack of Planning (UPPS_lack_planing), Sensation Seeking (UPPS_sensation_seeking), Positive Urgency (UPPS_positive_urgency), Negative Urgency (UPPS_negative_urgency) and Lack of Perserverance (UPPS_lack_perserverance) were included in the analysis.

A modified version (9) of Behavioral Inhibition & Behavioral Activation Scales (BIS/BAS)(10) reported by the participants was used to assess the psychobiological models of two broad motivational systems, the behavioral activation system (BAS) and the behavioral inhibition system (BIS). The BIS/BAS scale has a dimensional scale to assess Behavioral Inhibition (BISBAS_bis_sum). and three scales to assess Behavioral Activation: Drive (BISBAS_drive), Reward Responsiveness (BISBAS_reward_resp), and Fun Seeking (BISBAS_fun_seek). All the 4 scales were included in the current analysis.

Youth-report Prodromal Questionnaire Brief Version (PQ-B) (11) was used to assess the psychosis risk symptoms. The severity score of prodromal psychosis (PQB_psychosis_severity) was included in the current analysis.

Measurements and abbreviations of dimensional psychopathology related structures are listed in Table S3.

**Clinical Diagnoses of Psychiatric Disorders**

The categorical diagnostic assessment was measured with the computerized Kiddie Schedule for Affective Disorders and Schizophrenia (K-SADS) for DSM-5 (KSADS-5) (8,12). In the baseline assessment, modules covering externalizing, psychosis, and eating disorders were reported by parents, while modules of mood disorders, separation anxiety, social anxiety, generalized anxiety, sleep, and suicidality were reported by the participants with support from trained research staff. The rationale has been discussed in Barch et al., 2018 (8).

**Image Acquisition and Processing**

In the ABCD data study, MRI data of the participants were collected from 21 sites across the US. Structural and functional images were acquired using three 3 tesla scanner platforms: Siemens Prisma and Prisma Fit, General Electric Discovery MR750 and SIGNA Creator, and Phillips Medical System Ingenia and Achieva dStream; all were equipped with multi-channel coils in adult-size that are capable of multiband echo-planar imaging (EPI) acquisitions. Structural MRI images were collected with T1-weighted magnetization-prepared rapid acquisition gradient echo and T2-

weighted fast spin echo with variable flip angle scans. Resting-state and task functional images were collected with high spatial and temporal resolution simultaneous multi-slice /multiband EPI. The resting-state fMRI acquisition included four 5-min scans, during which participants were instructed to keep their eyes open and passively watch a crosshair. Parameters and the protocol of acquisition are detailed elsewhere (2).

Structural and functional MR images were preprocessed and housed in the ABCD-BIDS Community Collection (ABCC) from the Developmental Cognition and Neuroimaging (DCAN) Labs. The preprocessing abcd-hcp-pipeline mainly consists of two stages. The first stage is a modified version of Human Connectome Project (HCP)'s minimal preprocessing pipeline (13) to map fMRI volumes from the native space of each individual into a standard grayordinate spatial coordinate system, a combined cortical surface and subcortical volume coordinate system. The second stage used the DCAN BOLD Processing (DBP) software (14) to remove nuisance signals including artifacts induced by in-scanner head motion and participant respiration.

The modified HCP’s minimal preprocessing pipeline includes processing of both structural and functional MR images. For the structural MRIs, the processing includes denoising and N4 bias field correction, segmentation of brain tissue, volume-based spatial normalization of the sub-cortical brain and folding-based surface spatial normalization of the cortex. For fMRI, the processing includes correction of the gradient-nonlinearity-induced distortions, frame alignment, calculation of the motion parameters, non-linear registration to the MNI standard space, and volume-to-surface mapping to the grayordinate space.

The DBP is modified from the procedure of Power et al., 2014 (15), with correction of respiratory effects from motion estimates within multi-band data (14). The processing includes de-meaning and de-trending; nuisance-signal regression including mean time series from white matter and CSF; the global signal (GS); and the 24-parameter Volterra expansion from motion measures estimated by re-alignment (16,17); band-pass filtering between 0.008 and 0.09 Hz using a 2nd order Butterworth filter; and motion censoring. In the DBP the framewise displacement (FD) used for the motion censoring is corrected by filtering out the respiratory frequencies (0.31 Hz and 0.43 Hz). In the current analysis, frames of fMRI data with corrected FD > 0.2 mm were censored (14,15) before entering the construction of functional connectome. To ensure a reliable estimation of the resting-state functional connectivity (rsFC), 1645 participants with remaining time series < 6 min after censoring were excluded for further analysis (18).

**Construction of Functional Connectome**

Functional connectome was constructed from inter-regional BOLD synchrony of 352 areas across an individual's brain. For the cortex, the Gordon atlas (19) was used to divide the brain into 333 areas. The Gordon parcellation uses rsFC-boundary mapping to parcellate the cortex of the human brain, ensuring that vertices within the same parcel show homogenous rsFC patterns. The 333 cortical parcels were further assigned into 12 functional communities or networks using an Infomap based procedure (20). Because the identification of functional communities varies across studies due to the variance in methodological details, in order to facilitate comparison to other studies, we reassigned the original 12 functional networks of Gordon atlas (19) into the 7 network scheme (21) following the guideline of Uddin et.al. 2019 (22). The 7 functional networks include the Somatomotor Network (SMT), Visual Network (VIS), Default Mode Network (DMN), Frontoparietal Network (FPT), Salience/Ventral Attention Network (SAL/VAT), Dorsal Attention

Network (DAT), and Limbic Network (LIMB). 19 subcortical nuclei of left and right (L/R) Cerebellum, L/R Pallidum, L/R Caudate, L/R Putamen, L/R Accumbens, L/R Thalamus, L/R Amygdala, L/R Hippocampus, L/R Ventral Diencephalon, and Brain_Stem are assigned into the one Subcortical Network (SUB).

For each brain, the preprocessed BOLD signals within each parcel were extracted and averaged. The rsFC between every pair of parcels was measured as the Fisher Z-transformed Pearson's correlation between the two averaged time series. A 352 by 352 correlation matrix for each participant was thus obtained, which represents that brain's functional connectome profile.

**Covariates**

In the current study, we considered two tiers of covariates that may affect the CCA analysis (23). The Tier-1 covariates included the scanner type, head motion measured with mean framewise displacement (mean FD), and the number of retained frames after motion censoring. These Tier-1 covariates could induce artificial rsFC values, and the head motion during scanning could be associated with behavioral measures. Therefore, they might bias the CCA results with a spurious brain-behavior association. To eliminate such potential effects, we regressed out Tier-1 covariates from both brain and behavioral data in all our CCA analyses.

The Tier-2 covariates included 5 major sociodemographic variables, i.e., sex, age, race/ethnicity, household income, and parental education. These variables may have effects on the brain-behavior association, however a causal mechanism is unspecified. These variables might also proxy for both confounding factors and mediators or colliders simultaneously (23). Therefore, we conducted our original analyses without considering these variables and then conducted subsequent CCA analyses regressing out the Tier-2 covariates one at a time in addition to Tier-1 covariates, in order to investigate the impact of these Tier-2 covariates on the identified brain-behavior association.

**CCA within a Multiple Hold-out Framework**

The CCA analysis was performed within a nested 5-fold hold-out framework (Figure S2). The whole data set of 7382 participants was randomly split into a training set of 5906 participants (80% of the total samples) and a test set of 1476 participants (20% of the total samples). CCA model was trained in the training set and applied on the test set. Within the training set, a nested 5-fold hold out validation (80% vs 20% for each fold) was performed to tune the one hyperparameter, the number of principle components retained after the dimension reduction procedure (Figure S3), see bellowing for PCA-based dimension reduction. Note that, in the cross-validation, PCA was only performed on the training dataset, and then the resultant coefficients were applied to the test dataset.

**Pre-CCA Processing**

Before submitting them for CCA analysis, we performed two pre-CCA processing steps on the data sets.

The first was to eliminate spurious associations between the brain and behavior set introduced by data collection and head motion. The three Tier-I covariates: scanner type, head motion, and the number of retained frame were estimated and regressed from the both data sets (connectome and behaviors), and the residuals were transformed into a lower dimensional basis where data are exchangeable (24,25).

Second, the number of dimensions of the brain and behavioral data sets were separately reduced using principal component analysis (PCA). For the behavior set, 49 PCs were retained to keep all of the variance after residualizing. For the brain set, the number of PCs was set as an hyperparameter to be tuned within the nested 5-fold hold-out validation; the number of retained PCs was optimized to give the highest out-of-sample correlation. The searching space for PC number ranged from 49 (the rank of behavior data set) to 2000 (explaining ~80% variance of the brain data set), with a step of 50. Note that CCA is invariant to linear transformations, e.g. PCA in the current case, applied to either variable (24). That is, PCA does not change the result of CCA applied on the original feature space. However, removing the low-ranked PCs prevents small perturbations in the original data from causing instability in the CCA solutions and therefore overfitting.

**Canonical Correlation Analysis**

CCA is an analysis approach used to find multivariate associations between two data sets. After residualization and dimension reduction, the brain and behavior sets are full rank and noted as X and Y. CCA projects each set into K-dimensional latent spaces $\boldsymbol{U} = \boldsymbol{Y} \cdot \boldsymbol{A}$ and $\boldsymbol{V} = \boldsymbol{X} \cdot \boldsymbol{B}$, such that the correlations between each pair of canonical variates $\boldsymbol{u_k}$ and $\boldsymbol{v_k}$ are maximized, while canonical variates from the same set are orthogonal, i.e. $\boldsymbol{U^T U} = \boldsymbol{I}$ and $\boldsymbol{V}^T\boldsymbol{V} = \boldsymbol{I}$. Each pair of canonical variates $\boldsymbol{u_k}$ and $\boldsymbol{v_k}$ is called a mode, representing the latent dimensions linking the two data sets.

The number of modes is determined by the minimal rank of the two data sets, i.e. K = min(rank(Y), rank(X)). In the current case, K was 49, which is given by the rank of the behavioral set.

**Statistical Evaluation of the Canonical Variates**

We tested the statistical significances of the CCA modes within the training set. The test was performed using the permutation interference procedure of Winkler et al (24). In Winkler's procedure (24), the residualized datasets were projected to a lower dimensional basis so that the data were exchangeable. Then, in a stepwise manner, the significance of each pair of the canonical variates were inferred through permutation tests, while the variance explained by previous modes were removed from the data sets. The significance level was set at alpha = 0.05 from the 1000 permutations. The significance level was corrected using family wise error rate (FWER) correction, with a closed testing procedure.

For the 49 CCA modes, we further tested their out-of-sample generalizability. The assessment and statistical inference procedure are described previously (26,27). For each pair of canonical variates

$$\boldsymbol{u}_k = \boldsymbol{Y} \cdot \boldsymbol{a}_k$$
$$\boldsymbol{v}_k = \boldsymbol{X} \cdot \boldsymbol{b}_k$$

estimated in the training set, canonical weights were applied to the hold-out test set $\boldsymbol{Y}^*$ and $\boldsymbol{X}^*$, and the generalizability of mode $\boldsymbol{k}$ was assessed by the correlation between the projections

$$\rho = corr(\boldsymbol{Y}^* \boldsymbol{a}_k, X^* \boldsymbol{b}_k).$$

The null distribution of generalizability was constructed by

$$\rho_b = corr(\boldsymbol{X}^* \boldsymbol{a}'_k, \boldsymbol{Y}^* \boldsymbol{b}'_k),$$

where $\boldsymbol{a}'_k$ and $\boldsymbol{b}'_k$ were estimated by CCA performed on the permutated training set. The test for generalizability was performed in a stepwise manner. Matrix deflation with orthogonal projection was used to remove the variance explained by previous modes. The significance level was set at alpha = 0.05 from 1000 permutations. The significance level was corrected using family wise error rate (FWER) correction, with a closed testing procedure.

Finally, to evaluate the adequacy of brain-behavior prediction (28) for each mode, we examined the asymmetric redundancy index (29), which gives the mean variance of the behavioral data explained by the brain data set through a given mode. The normalized redundancy indicated the proportional contribution of each mode to the total variance of one of the two data sets that was explained by the other. It revealed an unequal pattern such that top-ranked CCA modes dominated the redundant variance. We conducted a permutation test for the normalized redundancy. Specifically, the CCA procedure was performed on a permutated brain and behavioral set, then the normalized brain and behavioral redundancy were calculated for each mode to generate the null distribution. The significance level was set at alpha = 0.05 from 1000 permutations. The significance level was corrected using family wise error rate (FWER) correction, with a closed testing procedure.

Table S4 summarizes the result of statistic tests for the significant CCA modes. Figure S4 and S5 show the behavioral and connectome loadings of the CCA modes 2-5.

**Predicting Behavioral Assessments from Mode-1 Connectome Variate**

CCA model was estimated from the training set following procedures described above. The first column vector $\boldsymbol{a}_1$ from the coefficient matrix $\boldsymbol{A}$ was used as the brain model. We then applied $\boldsymbol{a}_1$ connectome features in the test set to estimate the rsFC variate score in the unseen population $\boldsymbol{v}_1^* = X^*\boldsymbol{a}_1$. Finally, in the test set, Pearson's correlation tests were performed between the predicted brain variate score $\boldsymbol{v}_1^*$ and estimate and each of the 51 assessments in the behavior set $\boldsymbol{Y}^*$. The level of significance was at p = 0.05, multiple comparisons adjusted using the Benjamini and Hochberg procedure(30) to control the false discovery rate (FDR). The Pearson's correlation test and FDR adjustment were performed using the R package 'stats' v3.6.2.

**Association Between Connectome Variate, Clinical Diagnoses, and Status Transitions**

With the goal of determining the clinical utility of the connectome variate in predicting psychiatric disorders, we characterized associations between the connectome variate at baseline and clinical diagnoses at baseline and 2-year-follow-up. Towards this, we took a multi-pronged analytic approach that enabled us to investigate disorder-specific and transdiagnostic relationships and transitions between healthy and disordered states and between healthy and diagnosed states across different time-points. <u>To examine disorder-specific relationships</u>: All the 7382 participants with psychiatric disorders were grouped by their K-DSADS diagnoses, and the connectome variate score of each diagnostic group was compared with the group with no current psychiatric diagnosis using Welch two-sample t-tests. <u>To examine transdiagnostic relationships</u>, three analyses were conducted: First, we collapsed across diagnostic categories, grouping all participants by cumulative number of psychiatric disorders (0, 1, 2, or ≥3 current psychiatric diagnoses), and examining associations between the connectome variate and cumulative number of diagnoses at baseline and 2-year-follow-up. For this analysis, Welch two-sample t-tests (55) were used to compare differences between groups. Next, we stratified based upon connectome variate scores and examined the risk for developing one or more current psychiatric disorders in the total sample. This was done by stratifying and making group comparisons on psychiatric outcomes between participants with rsFC score above MEAN+1SD vs. under MEAN-1SD. Diagnostic Status Transitions: Lastly, we used data on current psychiatric disorders at baseline and 2-year-follow-up and the stability/persistence or change in status over time to stratify participants into four 'transition' group categories (Healthy-Persistent, Disorder-New-onset, Disorder-Remitted, and Disorder-Persistent) (see Fig. 8A). Diagnostic Status Transition Groups: The Healthy-persistent group (n=4699) was defined as participants who had 0 current psychiatric diagnoses at baseline and at 2-year-follow-up. This group was persistently mentally healthy and without psychiatric disorders over the study time period. They experienced no status transition over the study period. The Disorder-New-onset group (n=588) was defined as participants who had 0 current psychiatric diagnoses at baseline and ≥1 psychiatric diagnoses at 2-year-follow-up. This group went from having no psychiatric disorders to having one or more psychiatric disorders over the study time period. They experienced a status transition from healthy to diagnosed status. The Disorder-remitted group (n=787) was defined as participants who had ≥1 psychiatric diagnoses at baseline and who had 0 diagnoses at 2-year-follow-up. This group went from meeting criteria for one or more psychiatric disorders at baseline to not meeting criteria for any psychiatric disorders at the follow-up time point. They experienced a status transition from diagnosed to healthy status during the study period. The Disorder-persistent group (n=477) was defined as participants who had ≥1 psychiatric diagnoses at both baseline and 2-year-follow-up timepoints. This group had one or more psychiatric disorders at baseline and also at the follow-up timepoint, and thus experienced no status transition during the study period. Diagnostic Status Transitions Analysis: Having identified ‘transition’ groups, we then used ANOVAs to compare connectome variate scores across transition groups while adjusting for cumulative number of baseline psychiatric diagnoses (see Fig 8B).

### *Supplementary Results*

#### Inequality in the Spatial Distribution of rsFC Loadings for the Mode-1 Connectome Variate

The contribution of each rsFC to the composite brain score was measured by the rsFC loading, defined as the Pearson's correlation between the rsFC and the brain connectome variate across the training population. Here, as in the main manuscript, we only investigated the rsFC loadings of the first mode. A positive or a negative loading indicates that the given rsFC increases or decreases along the variate respectively. We define an ROI loading index by summing the loading of all the rsFCs from the given ROI; i.e. summing along each row in the loading matrix, which indicates the contribution of each brain ROI.

The Gini coefficient (29) was calculated to quantify the inequality across the 352 ROI sums (Figures S8C and G). Inequality of ROI loadings was tested for the positive and negative loading matrices separately. For each of the two loading matrices (Figures S6A and E), absolute rsFC loadings were summed at each ROI (Figures S6 B and F). Here absolute value was used for negative loadings because Gini coefficient only applies for non-negative values. We then permuted the elements in the loading matrix 1000 times and calculated the Gini coefficient from each of the permuted matrices forming the null distribution (Figures S6D and H). The p-value of the test was calculated by the proportion of permuted Gini coefficients exceeding the true one.

The Gini coefficient ranges between 0 to 1. A Gini of 0 indicates all the observed measurements are perfectly equal, while a Gini approaching 1 indicates all the measurements but one has a zero value, a completely unequal distribution. The calculation of the Gini coefficient and the related Lorenz curve (30) was using the R package 'DescTools' version 0.99.

(https://andrisignorell.github.io/DescTools/ ); see Cowell et al., 2000 (31) for further details on the Gini coefficient and the Lorenz curve.

**Cross-Domain Consistency of the Mode-1 Connectome Variate**

The rsFC pattern associated with the first CCA mode covaried with behavioral variates that correlated positively with cognitive functions and negatively with psychopathology related constructs. Such a transdiagnostic association implies that deficits of cognitive functions and severity of psychopathological problems may share a common set of neural processes. Alternatively, such an association may be driven by only one behavioral domain. For example, the first CCA mode may solely reflect an association between cognitive functions and rsFC. In that case, psychopathological measures may still show negative loadings due to the negative correlation between p- and g- factors.

To exclude this alternative hypothesis, we tested the cross-domain consistency of the CCA modes. Specifically, the same CCA procedure described above was conducted twice, but with behavioral assessments from only a single domain, cognition or psychopathology, at a time. In both cases, the first mode survived from both tests of within-sample significance and out-of-sample generalizability. Although the canonical correlation values differed between the CCA models estimated from cognition, psychopathology (Figure S7) or their combination (Figure 2), the connectome variates showed very high consistency among the three cases (combined vs. cognition only, $r = 0.990$; combined vs. psychopathology only, $r = 0.952$; Figure S7). Note that cross-domain consistency with high correlation (i.e. $|r| > 0.9$) was found to be specific to the first mode (see Table S5 for other significant modes).

These results lend support to the transdiagnostic nature of the connectome variate of the first mode and suggests that there may be a core neural substrate shared by both cognitive processes and psychopathology in this cohort.

**Impact of Sociodemographic Covariates on the first CCA mode**

We conducted sensitivity tests to evaluate how the first CCA mode was affected by sociodemographic factors. We focused on key sociodemographic variables (48), namely sex, race/ethnicity, household income, and parental education. We regressed out these sociodemographic covariates one at a time, in addition to the Tier-1 covariates, i.e. scanner type, mean frame displacement and remaining number of frames after censoring. As shown in Figure S8, the first CCA mode was reproduced in all cases, with a generalized canonical correlation ($\rho$ = 0.48~0.58) and similar connectome variate loading (r = 0.86~0.99), behavioral loading (r = 0.98~0.99) comparing with Figure 3.

**Connectome-based Prediction with CCA Modes Trained from Single-Domain Behaviors**

The connectome-based prediction in Figure 4 was replicated, but with CCA models trained with only one domain of behaviors, cognition or psychopathology. The weight matrix was applied to rsFC features in the hold-out test set and the predicted connectome variate score was correlated to the behaviors. As showing in Figure S9 and Figure S10 CCA models trained from each single-domain behavior set gives similar prediction performance as CCA model trained from the combined behavioral set (Figure 4).

*Supplementary Figures and Tables*

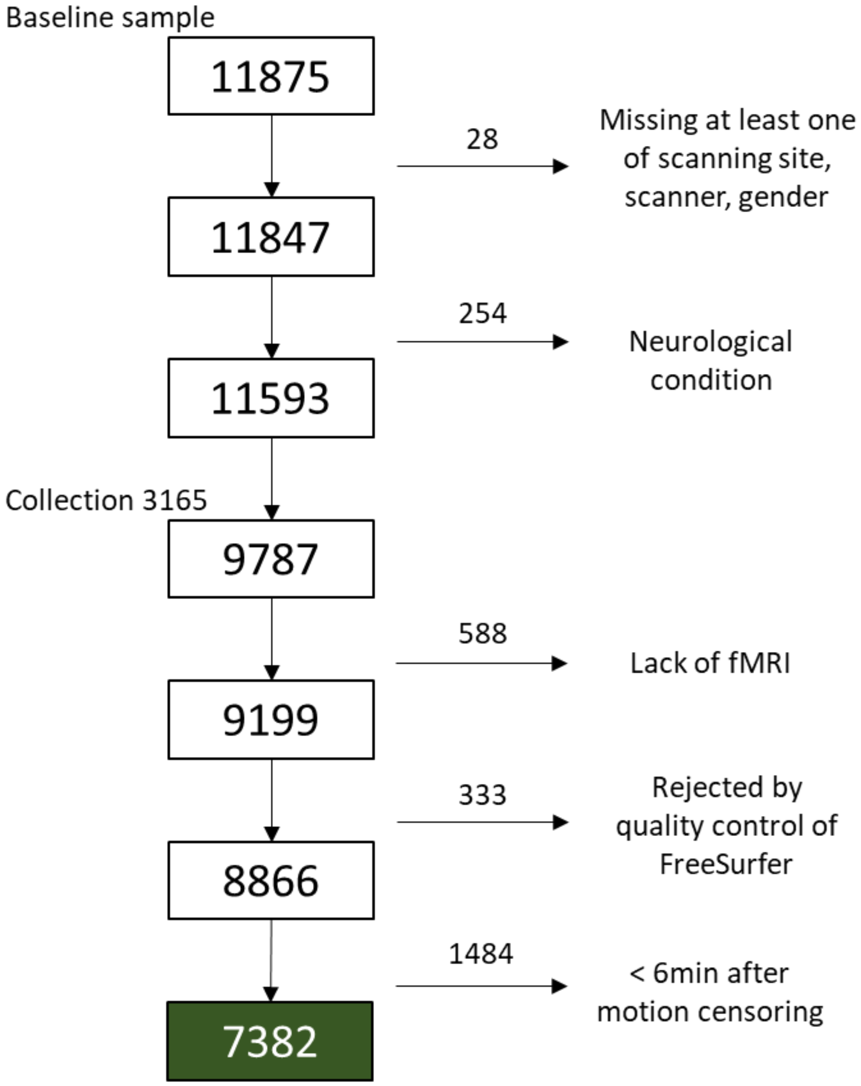


**Figure S1. Criteria for data exclusion used in this study.**

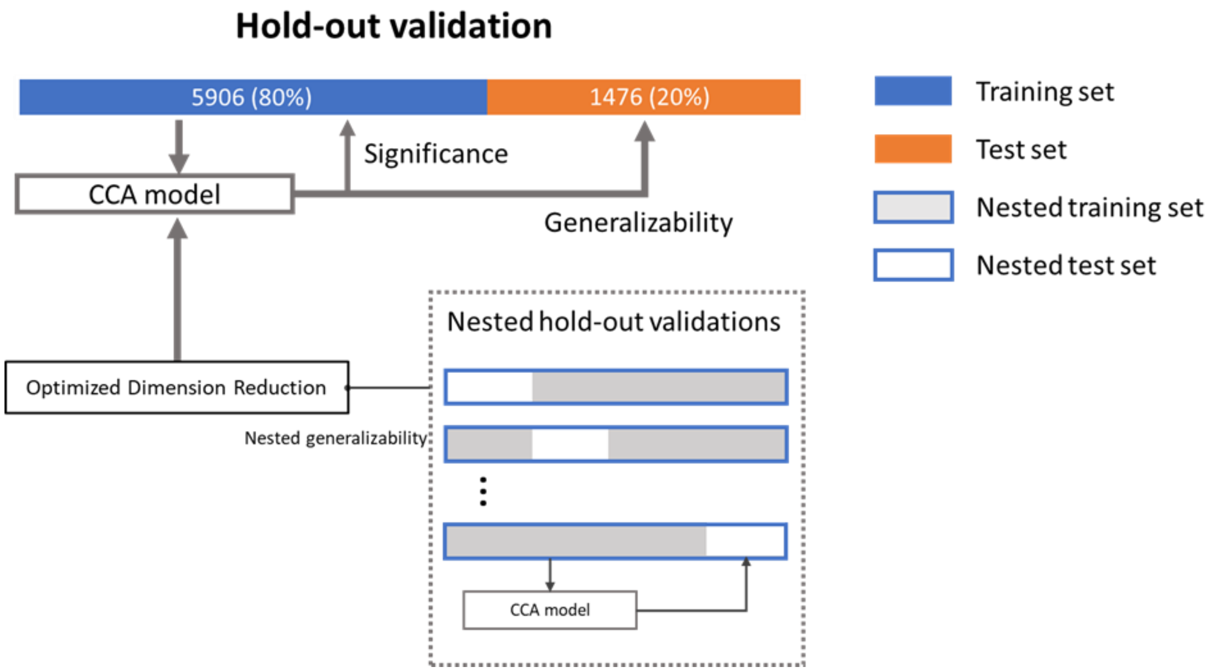


**Figure S2 Multiple hold-out framework for performing CCA analysis.**

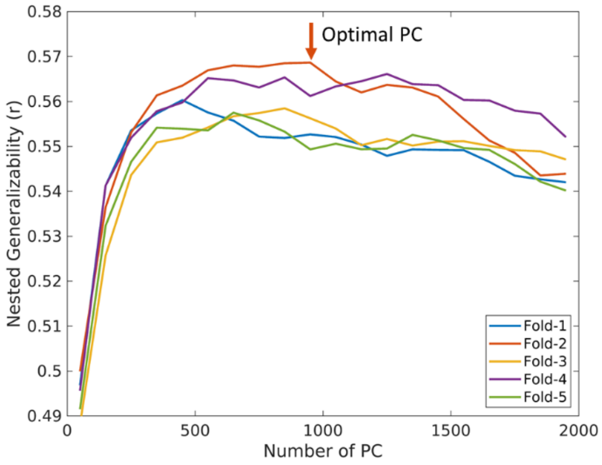


**Figure S3 Optimization of PCA-based dimension reduction in nested cross validation.** Line plots indicate the nested generalizability (averaged by each nested fold) along the number of retained PCs. The red arrow indicates the optimized PC for fold 2.

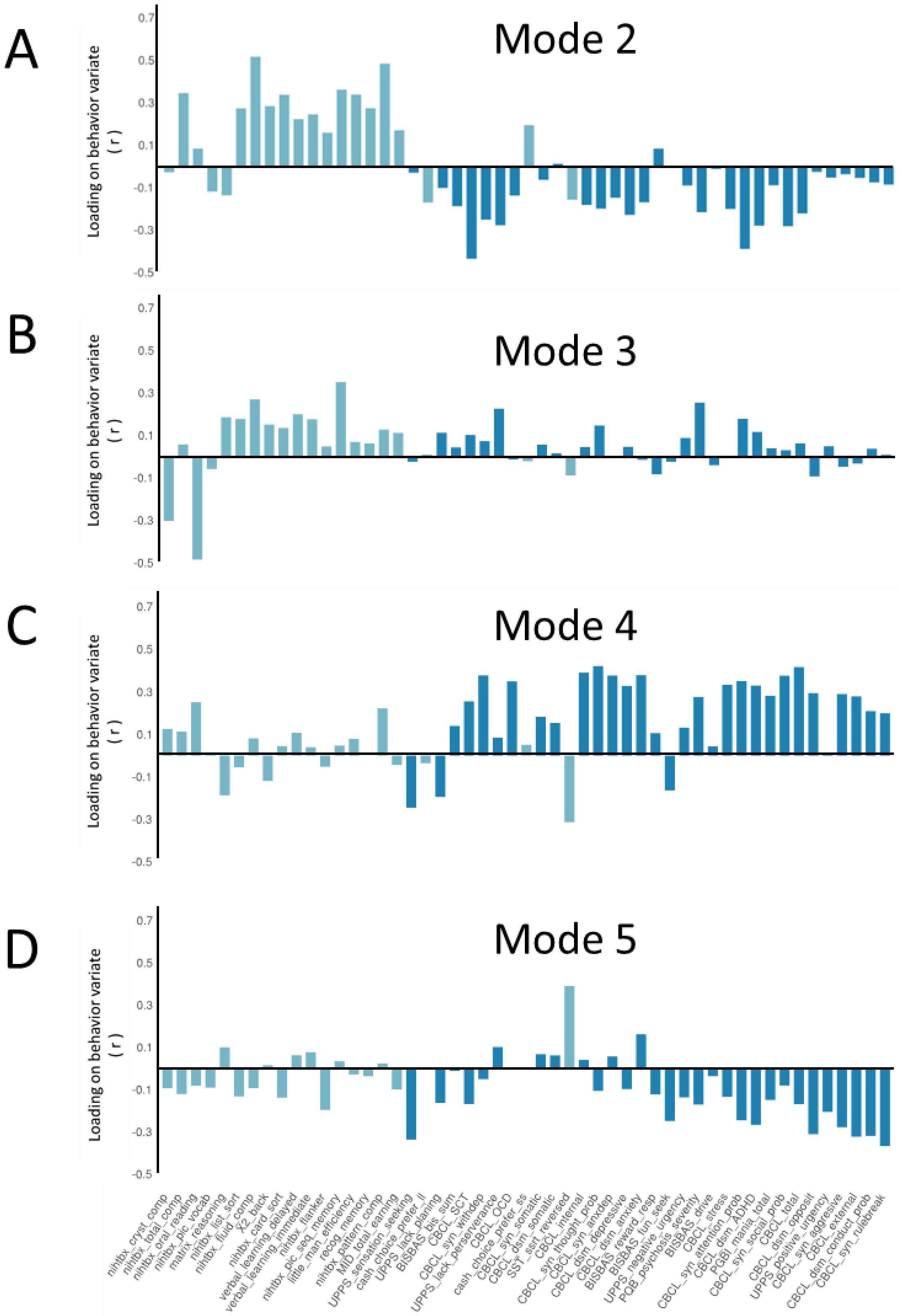


**Figure S4 Behavioral loadings of other significant modes. (A) Behavioral loadings of Mode 2. (B) Behavioral loadings of Mode 3. (C) Behavioral loadings of Mode 4. (D) Behavioral loadings of Mode 5. For visualization purpose, behavioral loadings are ordered as Figure 3A.**

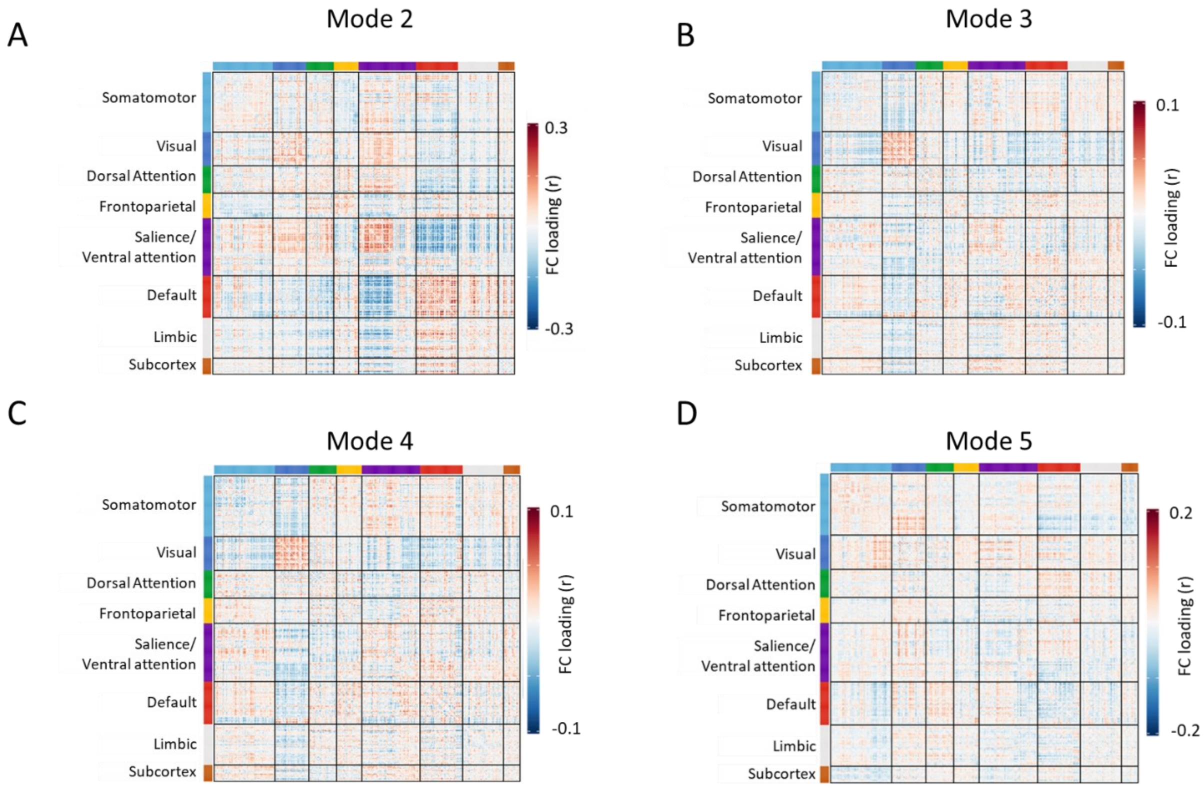


**Figure S5 RSFC loadings of the other significant modes. (A) RSFC loadings of Mode 2. (B) RSFC loadings of Mode 3. (C) RSFC loadings of Mode 4. (D) RSFC loadings of Mode 5.**

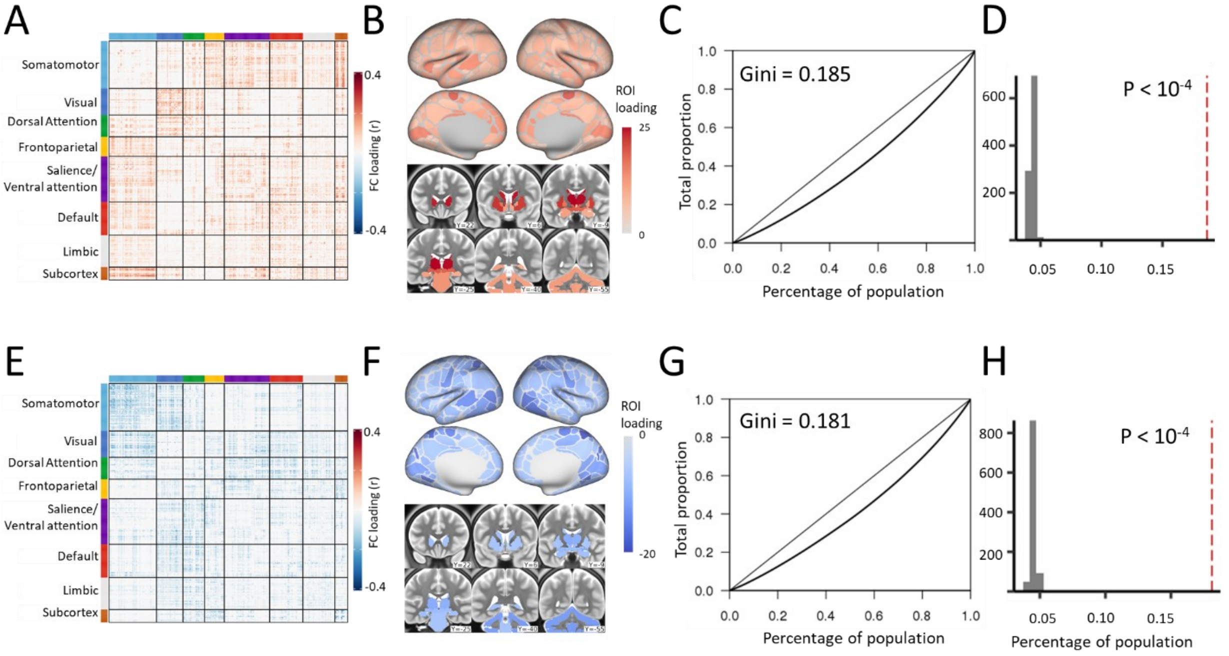


**Figure S6. (A) Matrix of positive resting-state functional connectivity (rsFC) loadings. (B) Sum of positive rsFC loadings by ROI. (C) Lorenz curve from the ROI sum of positive rsFC loadings. (D) Null distribution of Gini generated from permutated matrices of positive rsFC loadings. (E) Matrix of negative rsFC loadings. (F) Sum of negative rsFC loadings by ROI. (G) Lorenz curve from the ROI sum of negative rsFC loadings. (H) Null distribution of Gini generated from permutated matrices of negative rsFC loadings.**

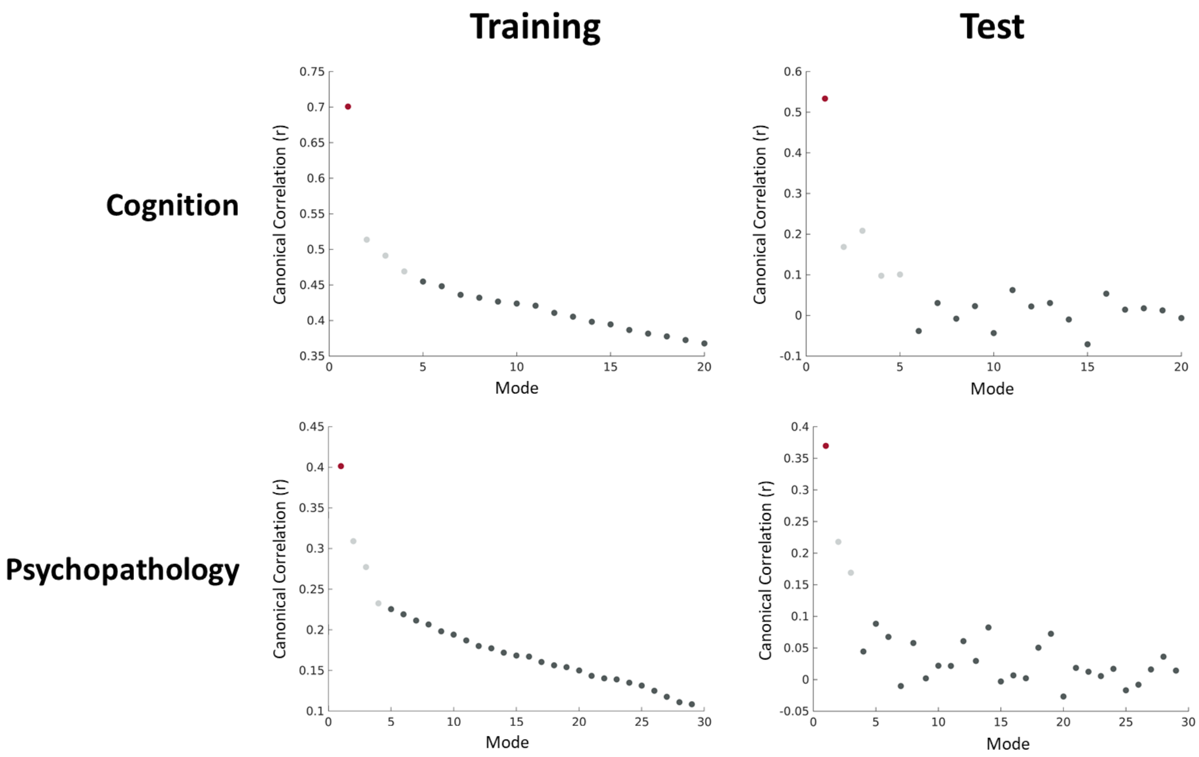


**Figure S7. Modes of canonical correlations when CCA was conducted with only one domain of behaviors.**

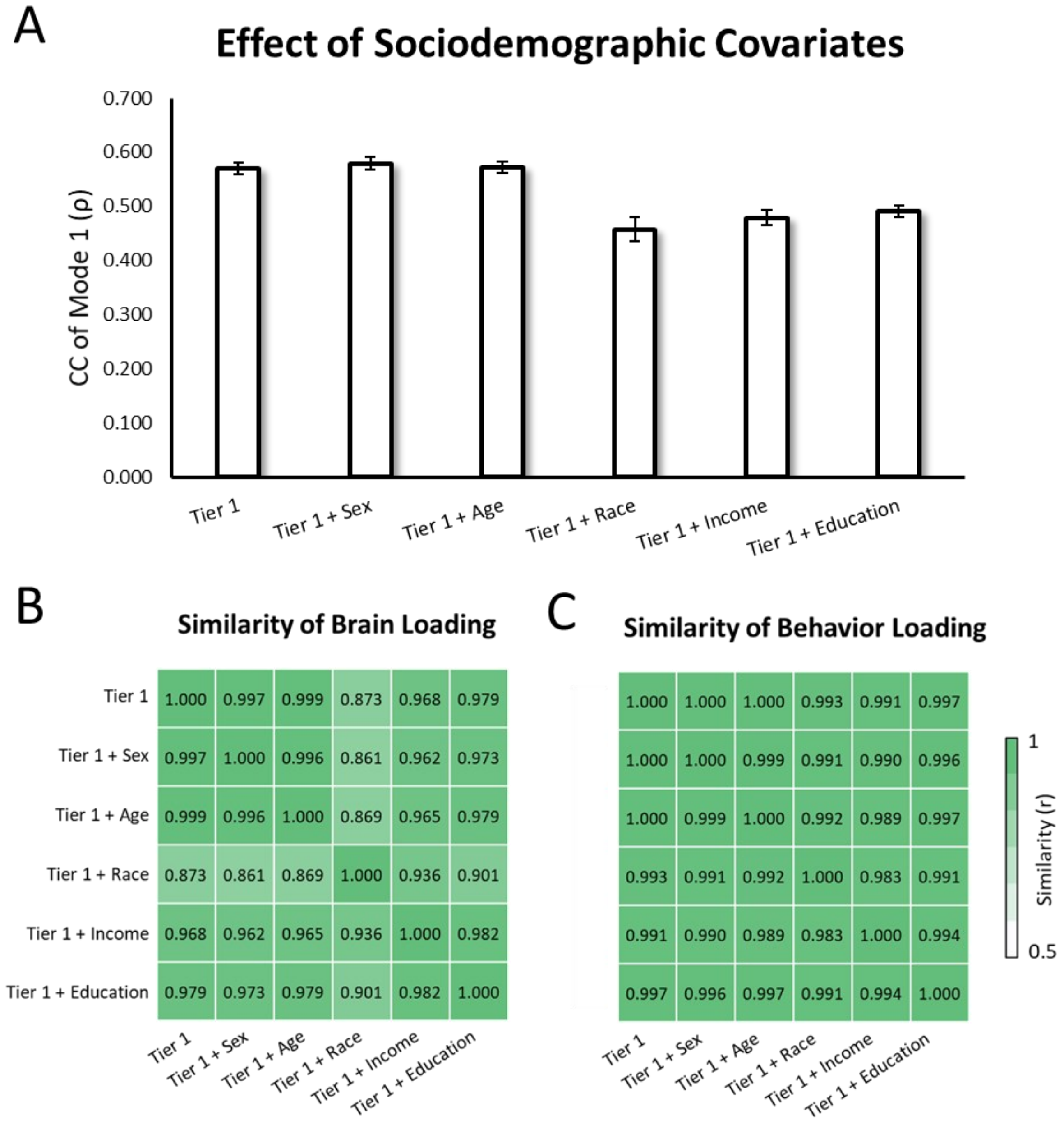


**Figure S8 Effect of sociodemographic covariates. (A) Generalized CC (B) Similarity of rsFC loading and (C) Similarity of behavior loading when sociodemographic variables were regressed out in addition to Tier-1 covariates.**

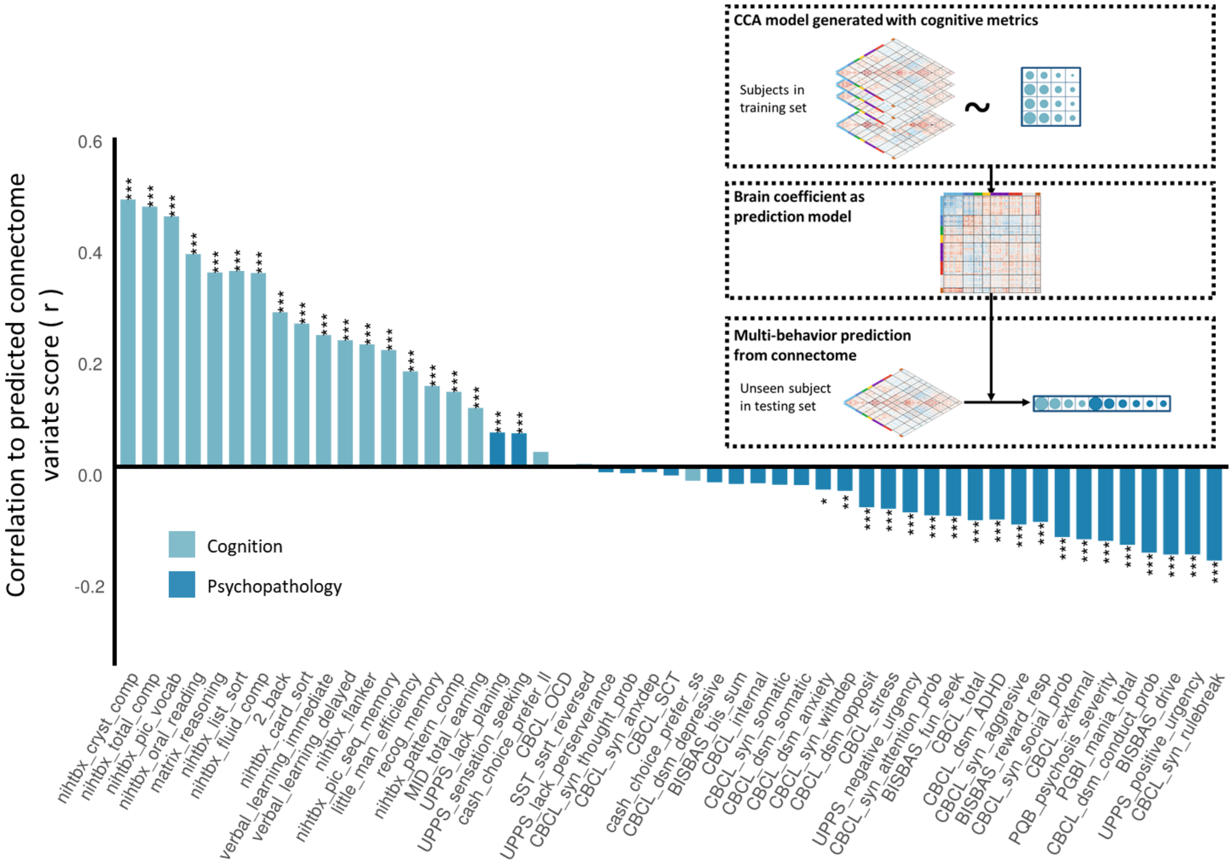


**Figure S9. Behavioral assessment prediction from the connectome variate score estimated from cognitive assessments.**

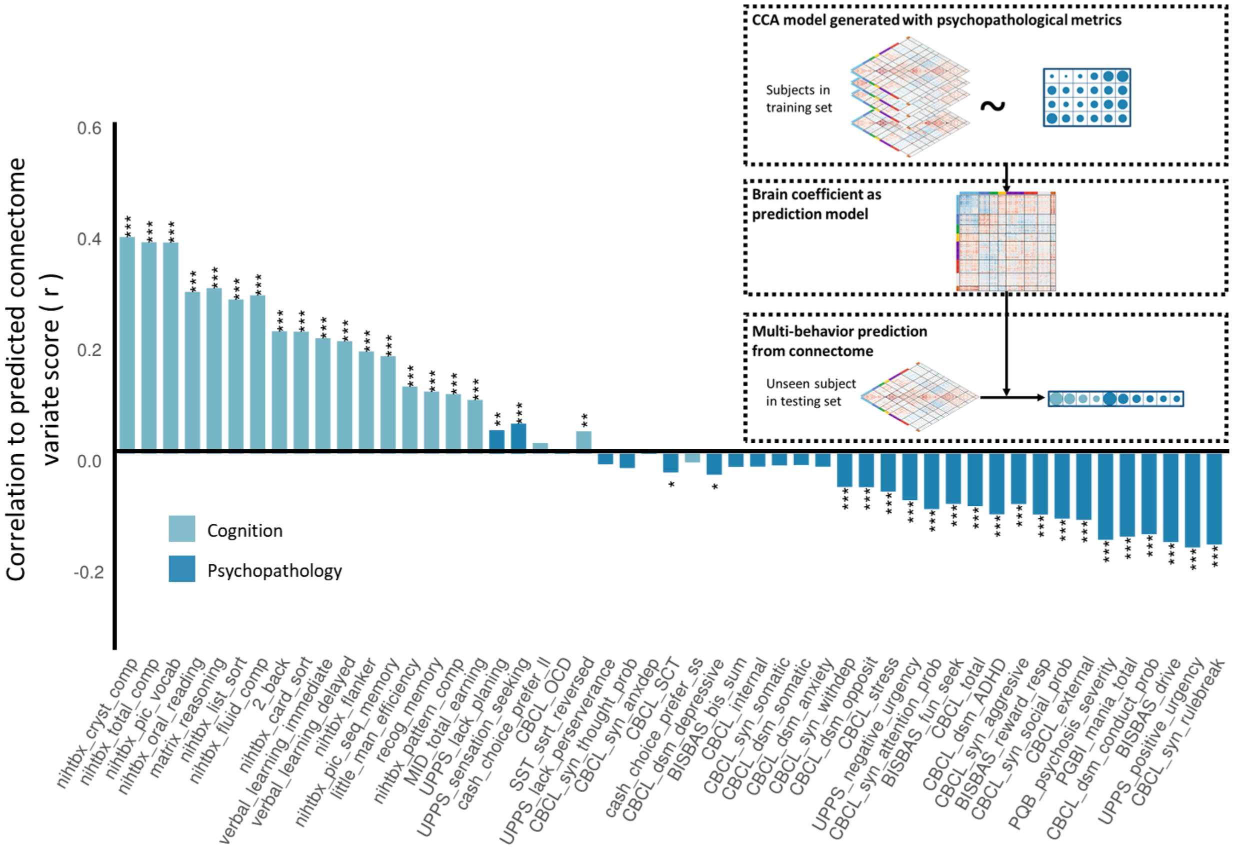


**Figure S10. Behavioral assessment prediction from the connectome variate score estimated from psychopathology related constructs.**

**Table S1. Demographic information of participants used in this study.**

| | |
|---|---|
| Age | 9.95 ± 0.62 y/o (Mean ± SD) |
| Gender | Female: 3714, Male: 3668 |
| Race | White: 4175<br>Hispanic: 1392<br>Black: 928<br>Unknown: 753<br>Asian: 134 |

**Table S2. Measurements and abbreviations for cognitive assessments used in this study.**

| Test | Measurement | Abbreviation |
|---|---|---|
| **Cognition Test:** | | |
| NIH Toolbox Flanker | Raw score of test | nihtbx_flanker |
| NIH Toolbox List Sorting Working Memory | Raw score of test | nihtbx_list_sort |
| NIH Toolbox Dimensional Change Card Sort | Raw score of test | nihtbx_card_sort |
| NIH Toolbox Oral Reading Recognition Test | Raw score of test | nihtbx_oral_reading |
| NIH Toolbox Pattern Comparison Processing Speed | Raw score of test | nihtbx_pattern_comp |
| NIH Toolbox Picture Sequence Memory Test | Raw score of test | nihtbx_pic_seq_memory |
| NIH Toolbox Picture Vocabulary Test | Raw score of test | nihtbx_pic_vocab |
| Fluid_intelligence | Raw composite score | nihtbx_fluidcomp |
| Crystal_intelligence | Raw composite score | nihtbx_cryst_comp |
| Total_intellegence | Raw composite score | nihtbx_total_comp |
| Rey Auditory Verbal Learning | Accuracy in the immediate test | verbal_learning_immediate |
| | Accuracy in the delayed test | verbal_learning_delayed |
| Cash Choice Task | Choice of short smaller-sooner | cash_choice_prefer_ss |
| | Choice of short larger-later | cash_choice_prefer_ll |
| Little Man Task | Efficiency ratio | little_man_efficiency |
| Matrix Reasoning Test | Total scaled score | matrix_reasoning |
| Recognition memory test | *d'* | recog_memory |
| **FMRI Tasks:** | | |
| Emotional n-back（2-back – 0-back） | % correct on 2-back - % correct on 0-back | 2_back |
| Stop-signal reaction time | -1*(stop – signal RT) | SST_ssrt_reversed |
| Monetary Incentive Delay | Mean earnings | MID_total_earning |

**Table S3. Measurements and abbreviations of dimensional psychopathological assessments used in this study.**

| Questionnaire | Measurement | Abbreviation |
|---|---|---|
| **Children Behavior Check List:** | | |
| Syndrome scales | Anxious/Depressive symptoms | CBCL_syn_anxdep |
| | Withdrawn/Depressed symptoms | CBCL_syn_withdep |
| | Somatic Complaints | CBCL_syn_somatic |
| | Social Problems | CBCL_syn_social_prob |
| | Thought Problems | CBCL_syn_thought_prob |
| | Attention Problems | CBCL_syn_attention_prob |
| | Rule-Breaking Behavior | CBCL_syn_rulebreak |
| | Aggressive Behavior | CBCL_syn_aggresive |
| | Internalizing Problems | CBCL_internal |
| | Externalizing Problems | CBCL_external |
| | Total Problems | CBCL_total |
| DSM-oriented scales | Depressive Problems | CBCL_dsm_depressive |
| | Anxiety Problems | CBCL_dsm_anxiety |
| | Somatic Problems | CBCL_dsm_somatic |
| | Attention-deficit/hyperactivity Disorder | CBCL_dsm_ADHD |
| | Oppositional Defiant Problems | CBCL_dsm_opposit |
| | Conduct Problems | CBCL_dsm_conduct_prob |
| 2007-Scale Scores | Sluggish Cognitive Tempo | CBCL_SCT |
| | Obsessive-Compulsive Problems | CBCL_OCD |
| | Stress Problems | CBCL_stress |
| **UPPS-P:** | Lack of Planning | UPPS_lack_planing |
| | Sensation Seeking | UPPS_sensation_seeking |
| | Positive Urgency | UPPS_positive_urgency |
| | Negative Urgency | UPPS_negative_urgency |
| | Lack of Perseverance | UPPS_lack_perserverance |
| **Behavioral Inhibition & Behavioral Activation Scales (BIS/BAS):** | | |
| | Drive | BISBAS_drive |
| | Reward responsiveness | BISBAS_reward_resp |
| | Fun seeking | BISBAS_fun_seek |
| | BIS total score | BISBAS_bis_total |
| **Prodromal Questionnaire Brief Version (PQ-B) :** | | |
| | Severity score of prodromal psychosis | PQB_psychosis_severity |
| **Ten-item Mania Scale:** | | |
| | Total score of mania | PGBI_mania_total |

**Table S4. Summary of the CCA analysis**

| Data split | Brain PC | Behavior PC | Correlation | Holdout correlation | Brain redundancy ($10^{-3}$) | Behavioral redundancy ($10^{-3}$) | Brain redundancy normalized (%) | Behavioral redundancy normalized (%) |
|---|---|---|---|---|---|---|---|---|
| **First mode of connectome-behavior association** | | | | | | | | |
| Fold 1 | 650 | 49 | 0.682$^{***}$ | 0.560$^{***}$ | 2.605$^{***}$ | 49.122$^{***}$ | 28.344$^{***}$ | 29.224$^{***}$ |
| **Fold 2** | **1050** | **49** | **0.725**$^{***}$ | **0.560**$^{***}$ | **2.653**$^{***}$ | **57.170**$^{***}$ | **26.442**$^{***}$ | **23.840**$^{***}$ |
| Fold 3 | 750 | 49 | 0.694$^{***}$ | 0.583$^{***}$ | 2.626$^{***}$ | 51.054$^{***}$ | 27.923$^{***}$ | 27.127$^{***}$ |
| Fold 4 | 950 | 49 | 0.715$^{***}$ | 0.566$^{***}$ | 2.647$^{***}$ | 56.250$^{***}$ | 26.323$^{***}$ | 25.281$^{***}$ |
| Fold 5 | 750 | 49 | 0.692$^{***}$ | 0.584$^{***}$ | 2.532$^{***}$ | 50.888$^{***}$ | 26.884$^{***}$ | 27.172$^{***}$ |
| **Second mode of connectome-behavior association** | | | | | | | | |
| Fold 1 | 650 | 49 | 0.480$^{***}$ | 0.207$^{***}$ | 0.420$^{***}$ | 9.252$^{***}$ | 4.574$^{***}$ | 5.504$^{*}$ |
| **Fold 2** | **1050** | **49** | **0.557**$^{***}$ | **0.174**$^{***}$ | **0.508**$^{***}$ | **17.328**$^{***}$ | **5.065**$^{***}$ | **7.226**$^{**}$ |
| Fold 3 | 750 | 49 | 0.500$^{***}$ | 0.233$^{***}$ | 0.584$^{***}$ | 14.054$^{***}$ | 6.204$^{***}$ | 7.467$^{*}$ |
| Fold 4 | 950 | 49 | 0.542$^{***}$ | 0.195$^{***}$ | 0.602$^{***}$ | 14.532$^{***}$ | 5.982$^{***}$ | 6.531$^{*}$ |
| Fold 5 | 750 | 49 | 0.505$^{***}$ | 0.206$^{***}$ | 0.639$^{***}$ | 15.082$^{***}$ | 6.781$^{***}$ | 8.053$^{**}$ |
| **Third mode of connectome-behavior association** | | | | | | | | |
| Fold 1 | 650 | 49 | 0.468$^{***}$ | 0.224$^{***}$ | 0.477$^{***}$ | 6.769$^{***}$ | 5.185$^{***}$ | 4.027 |
| **Fold 2** | **1050** | **49** | **0.542**$^{***}$ | **0.211**$^{***}$ | **0.383**$^{***}$ | **7.297**$^{***}$ | **3.818**$^{***}$ | **3.043** |
| Fold 3 | 750 | 49 | 0.484$^{***}$ | 0.195$^{***}$ | 0.332$^{***}$ | 4.452$^{***}$ | 3.532$^{**}$ | 2.366 |
| Fold 4 | 950 | 49 | 0.519$^{***}$ | 0.175$^{***}$ | 0.318$^{***}$ | 9.176$^{***}$ | 3.164$^{*}$ | 4.124 |
| Fold 5 | 750 | 49 | 0.488$^{***}$ | 0.176$^{***}$ | 0.379$^{***}$ | 4.603$^{***}$ | 4.020$^{***}$ | 2.458 |
| **Fourth mode of connectome-behavior association** | | | | | | | | |
| Fold 1 | 650 | 49 | 0.451$^{***}$ | 0.212$^{***}$ | 0.293$^{***}$ | 6.839$^{***}$ | 3.184$^{***}$ | 4.069 |
| **Fold 2** | **1050** | **49** | **0.536**$^{***}$ | **0.195**$^{***}$ | **0.362**$^{***}$ | **5.992**$^{***}$ | **3.609**$^{**}$ | **2.499** |
| Fold 3 | 750 | 49 | 0.479$^{***}$ | 0.194$^{***}$ | 0.279$^{***}$ | 7.588$^{***}$ | 2.963 | 4.032 |
| Fold 4 | 950 | 49 | 0.512$^{***}$ | 0.201$^{***}$ | 0.338$^{***}$ | 4.551$^{***}$ | 3.358$^{*}$ | 2.046 |
| Fold 5 | 750 | 49 | 0.473$^{***}$ | 0.165$^{***}$ | 0.304$^{***}$ | 6.304$^{***}$ | 3.231$^{*}$ | 3.366 |
| **Fifth mode of connectome-behavior association** | | | | | | | | |
| Fold 1 | 650 | 49 | 0.442$^{***}$ | 0.155$^{***}$ | 0.239$^{***}$ | 4.804$^{***}$ | 2.599 | 2.858 |
| **Fold 2** | **1050** | **49** | **0.516**$^{***}$ | **0.148**$^{***}$ | **0.314**$^{***}$ | **10.816**$^{***}$ | **3.134**$^{*}$ | **4.510** |
| Fold 3 | 750 | 49 | 0.460$^{***}$ | 0.195$^{***}$ | 0.277$^{***}$ | 8.175$^{***}$ | 2.947 | 4.344 |
| Fold 4 | 950 | 49 | 0.507$^{***}$ | 0.135$^{***}$ | 0.282$^{***}$ | 7.560$^{***}$ | 2.802 | 3.398 |
| Fold 5 | 750 | 49 | 0.470$^{***}$ | 0.208$^{***}$ | 0.233$^{***}$ | 7.526$^{***}$ | 2.473 | 4.019 |

Fold 2 was shown in the main text as a representative result

* $p_{fwer}<0.05$

** $p_{fwer}<0.01$

*** $p_{fwer}<0.001$

**Table S5 Cross-domain consistency for connectome variates of significant CCA modes**

| | Combined - Cognition (\|r\|) | Combined - Psychopathology (\|r\|) | Psychopathology - Cognition (\|r\|) |
|---|---|---|---|
| Mode 1 | 0.99 | 0.95 | 0.93 |
| Mode 2 | 0.62 | 0.80 | 0.28 |
| Mode 3 | 0.46 | 0.19 | 0.21 |
| Mode 4 | 0.43 | 0.23 | 0.17 |
| Mode 5 | 0.01 | 0.25 | 0.22 |

**Table S6 Prospective behavior prediction of the Mode-1 connectome variate**

| | Baseline | | | Year 2 | | |
|---|---|---|---|---|---|---|
| | r | p | p.fdr | r | p | p.fdr |
| nihtbx_cryst_comp | 0.4850 | 0.0000 | 0.0000 | 0.4868 | 0.0000 | 0.0000 |
| nihtbx_total_comp | 0.4674 | 0.0000 | 0.0000 | - | - | - |
| nihtbx_pic_vocab | 0.4573 | 0.0000 | 0.0000 | 0.4816 | 0.0000 | 0.0000 |
| nihtbx_oral_reading | 0.3837 | 0.0000 | 0.0000 | 0.3730 | 0.0000 | 0.0000 |
| matrix_reasoning | 0.3546 | 0.0000 | 0.0000 | - | - | - |
| nihtbx_list_sort | 0.3522 | 0.0000 | 0.0000 | - | - | - |
| nihtbx_fluid_comp | 0.3455 | 0.0000 | 0.0000 | - | - | - |
| 2_back | 0.2787 | 0.0000 | 0.0000 | 0.3347 | 0.0000 | 0.0000 |
| nihtbx_card_sort | 0.2557 | 0.0000 | 0.0000 | - | - | - |
| verbal_learning_immediate | 0.2447 | 0.0000 | 0.0000 | 0.2297 | 0.0000 | 0.0000 |
| verbal_learning_delayed | 0.2338 | 0.0000 | 0.0000 | 0.2217 | 0.0000 | 0.0000 |
| nihtbx_flanker | 0.2174 | 0.0000 | 0.0000 | 0.2469 | 0.0000 | 0.0000 |
| nihtbx_pic_seq_memory | 0.2090 | 0.0000 | 0.0000 | 0.2300 | 0.0000 | 0.0000 |
| little_man_efficiency | 0.1720 | 0.0000 | 0.0000 | - | - | - |
| recog_memory | 0.1459 | 0.0000 | 0.0000 | 0.1440 | 0.0000 | 0.0000 |
| nihtbx_pattern_comp | 0.1346 | 0.0000 | 0.0000 | 0.1423 | 0.0000 | 0.0000 |
| MID_total_earning | 0.1048 | 0.0000 | 0.0000 | 0.0556 | 0.0001 | 0.0012 |
| UPPS_lack_planing | 0.0647 | 0.0000 | 0.0000 | 0.0975 | 0.0000 | 0.0000 |
| UPPS_sensation_seeking | 0.0673 | 0.0000 | 0.0000 | 0.0165 | 0.1806 | 1.0000 |
| cash_choice_prefer_ll | 0.0305 | 0.0087 | 0.0959 | - | - | - |
| CBCL_OCD | 0.0070 | 0.5490 | 1.0000 | 0.0437 | 0.0055 | 0.0712 |
| SST_ssrt_reversed | 0.0101 | 0.3873 | 1.0000 | 0.0486 | 0.0001 | 0.0012 |
| UPPS_lack_perserverance | -0.0058 | 0.6200 | 1.0000 | -0.0667 | 0.0000 | 0.0000 |
| CBCL_syn_thought_prob | -0.0088 | 0.4513 | 1.0000 | 0.0209 | 0.0911 | 0.7285 |
| CBCL_syn_anxdep | -0.0069 | 0.5542 | 1.0000 | 0.0125 | 0.3135 | 1.0000 |
| CBCL_SCT | -0.0127 | 0.2747 | 1.0000 | 0.0116 | 0.3472 | 1.0000 |
| cash_choice_prefer_ss | -0.0246 | 0.0344 | 0.2824 | 0.0952 | 0.0000 | 0.0000 |
| CBCL_dsm_depressive | -0.0251 | 0.0314 | 0.2824 | - | - | - |
| BISBAS_bis_sum | -0.0228 | 0.0503 | 0.3518 | 0.0001 | 0.9944 | 1.0000 |
| CBCL_internal | -0.0267 | 0.0218 | 0.2176 | -0.0027 | 0.8251 | 1.0000 |
| CBCL_syn_somatic | -0.0311 | 0.0074 | 0.0894 | -0.0279 | 0.0240 | 0.2398 |
| CBCL_dsm_somatic | -0.0324 | 0.0054 | 0.0704 | -0.0258 | 0.0367 | 0.3304 |
| CBCL_dsm_anxiety | -0.0382 | 0.0010 | 0.0144 | -0.0083 | 0.5002 | 1.0000 |
| CBCL_syn_withdep | -0.0388 | 0.0009 | 0.0129 | -0.0125 | 0.3122 | 1.0000 |
| CBCL_dsm_opposit | -0.0724 | 0.0000 | 0.0000 | -0.0571 | 0.0000 | 0.0001 |
| CBCL_stress | -0.0727 | 0.0000 | 0.0000 | -0.0329 | 0.0078 | 0.0856 |
| UPPS_negative_urgency | -0.0779 | 0.0000 | 0.0000 | -0.0493 | 0.0001 | 0.0010 |
| CBCL_syn_attention_prob | -0.0852 | 0.0000 | 0.0000 | -0.0498 | 0.0001 | 0.0009 |
| BISBAS_fun_seek | -0.0885 | 0.0000 | 0.0000 | -0.0618 | 0.0000 | 0.0000 |
| CBCL_total | -0.0942 | 0.0000 | 0.0000 | -0.0552 | 0.0000 | 0.0001 |
| CBCL_dsm_ADHD | -0.0928 | 0.0000 | 0.0000 | -0.0558 | 0.0000 | 0.0001 |
| CBCL_syn_aggresive | -0.1014 | 0.0000 | 0.0000 | -0.0343 | 0.0055 | 0.0712 |
| BISBAS_reward_resp | -0.0983 | 0.0000 | 0.0000 | -0.0749 | 0.0000 | 0.0000 |
| CBCL_syn_social_prob | -0.1240 | 0.0000 | 0.0000 | -0.0709 | 0.0000 | 0.0000 |
| CBCL_external | -0.1289 | 0.0000 | 0.0000 | -0.0920 | 0.0000 | 0.0000 |
| PQB_psychosis_severity | -0.1355 | 0.0000 | 0.0000 | -0.1427 | 0.0000 | 0.0000 |
| PGBI_mania_total | -0.1409 | 0.0000 | 0.0000 | -0.1216 | 0.0000 | 0.0000 |
| CBCL_dsm_conduct_prob | -0.1527 | 0.0000 | 0.0000 | -0.1135 | 0.0000 | 0.0000 |
| BISBAS_drive | -0.1591 | 0.0000 | 0.0000 | -0.1256 | 0.0000 | 0.0000 |
| UPPS_positive_urgency | -0.1547 | 0.0000 | 0.0000 | -0.1157 | 0.0000 | 0.0000 |
| CBCL_syn_rulebreak | -0.1691 | 0.0000 | 0.0000 | -0.1106 | 0.0000 | 0.0000 |

**Table S7 Clinical Diagnosis as Function of Connectome Variate Score**

| Percentage of CVS* | Baseline | | | Year 2 | | |
|---|---|---|---|---|---|---|
| | No Diagnosis (N = 5927) | Any Diagnosis (N = 1454) | Total (N = 7381) | No Diagnosis (N = 5488) | Any Diagnosis (N = 3795) | Total (N = 6551) |
| High | 950 | 168 | 1158 | 850 | 142 | 992 |
| Middle | 4199 | 986 | 5105 | 3795 | 723 | 4518 |
| Low | 858 | 300 | 1118 | 843 | 198 | 1041 |

* CVS is short for connectome variate score.

Participants were grouped by the percentage of their CVS in the population.

High: ≥1SD above the Mean; Middle: within 1 SD of the Mean; Low: ≤ 1SD below the Mean.

### *References*


1. Luciana M, Bjork JM, Nagel BJ, Barch DM, Gonzalez R, Nixon SJ, Banich MT (2018): Adolescent neurocognitive development and impacts of substance use: Overview of the adolescent brain cognitive development (ABCD) baseline neurocognition battery. *Developmental Cognitive Neuroscience* 32: 67–79.
2. Casey BJ, Cannonier T, Conley MI, Cohen AO, Barch DM, Heitzeg MM, *et al.* (2018): The Adolescent Brain Cognitive Development (ABCD) study: Imaging acquisition across 21 sites. *Developmental Cognitive Neuroscience* 32: 43–54.
3. Rosenberg MD, Martinez SA, Rapuano KM, Conley MI, Cohen AO, Cornejo MD, *et al.* (2020): Behavioral and Neural Signatures of Working Memory in Childhood. *The Journal of Neuroscience* 40: 5090–5104.
4. Achenbach TM, Verhulst F (2010): *Achenbach System of Empirically Based Assessment (ASEBA)*. Burlington, Vermont.
5. Youngstrom EA, Frazier TW, Demeter C, Calabrese JR, Findling RL (2008): Developing a 10-Item Mania Scale from the Parent General Behavior Inventory for Children and Adolescents. *Journal of Clinical Psychiatry* 69: 831–839.
6. Youngstrom EA, Findling RL, Danielson CK, Calabrese JR (2001): Discriminative validity of parent report of hypomanic and depressive symptoms on the general behavior inventory. *Psychological Assessment* 13: 267–276.
7. Zapolski TCB, Stairs AM, Settles RF, Combs JL, Smith GT (2010): The measurement of dispositions to rash action in children. *Assessment* 17: 116–125.
8. Barch DM, Albaugh MD, Avenevoli S, Chang L, Clark DB, Glantz MD, *et al.* (2018): Demographic, physical and mental health assessments in the adolescent brain and cognitive development study: Rationale and description. *Developmental Cognitive Neuroscience* 32: 55–66.

9. Pagliaccio D, Luking KR, Anokhin AP, Gotlib IH, Hayden EP, Olino TM, *et al.* (2016): Revising the BIS/BAS Scale to study development: Measurement invariance and normative effects of age and sex from childhood through adulthood. *Psychological Assessment* 28: 429–442.
10. Carver CS, White TL (1994): Behavioral inhibition, behavioral activation, and affective responses to impending reward and punishment: The BIS/BAS Scales. *Journal of Personality and Social Psychology* 67: 319–333.
11. Loewy RL, Pearson R, Vinogradov S, Bearden CE, Cannon TD (2011): Psychosis risk screening with the Prodromal Questionnaire - Brief Version (PQ-B). *Schizophrenia Research* 129: 42–46.
12. Kobak KA, Kratochvil CJ, Stanger C, Kaufman J (2013): Computerized screening of comorbidity in adolescents with substance or psychiatric disorders. *Anxiety Disorders and Depression(La Jolaa, CA)*.
13. Glasser MF, Sotiropoulos SN, Wilson JA, Coalson TS, Fischl B, Andersson JL, *et al.* (2013): The minimal preprocessing pipelines for the Human Connectome Project. *NeuroImage* 80: 105–124.
14. Fair DA, Miranda-Dominguez O, Snyder AZ, Perrone A, Earl EA, Van AN, *et al.* (2020): Correction of respiratory artifacts in MRI head motion estimates. *NeuroImage* 208: 116400.
15. Power JD, Mitra A, Laumann TO, Snyder AZ, Schlaggar BL, Petersen SE (2014): Methods to detect, characterize, and remove motion artifact in resting state fMRI. *NeuroImage* 84: 320–341.
16. Yan CG, Cheung B, Kelly C, Colcombe S, Craddock RC, Di Martino A, *et al.* (2013): A comprehensive assessment of regional variation in the impact of head micromovements on functional connectomics. *NeuroImage* 76: 183–201.
17. Friston KJ, Williams S, Howard R, Frackowiak RSJ, Turner R (1996): Movement-related effects in fMRI time-series. *Magnetic Resonance in Medicine* 35: 346–355.
18. Birn RM, Molloy EK, Patriat R, Parker T, Meier TB, Kirk GR, *et al.* (2013): The effect of scan length on the reliability of resting-state fMRI connectivity estimates. *NeuroImage* 83: 550–558.
